# Methodology for a freshly engineered or cryo-preserved 3D tuberculoma bioplatform for studying tuberculosis biology and high-content screening of therapeutics

**DOI:** 10.1101/2025.08.27.672606

**Authors:** Suraj B. Sable, Allison Kline, Wen Li, James E. Posey

**Affiliations:** Division of Tuberculosis Elimination, National Center for HIV, Viral Hepatitis, STD, and TB Prevention, Centers for Disease Control and Prevention, Atlanta, GA, USA

**Author notes:** **Correspondence**: Suraj Sable.

**Keywords:** Tuberculosis, 3D cell culture, Granuloma model, Physiological microsystems, Shelf-stable, *M. marinum*, High-throughput screening, Host-directed therapy

## Abstract

Tuberculomas are the conglomeration of tuberculous granulomas into structurally organized three-dimensional (3D) masses that result from *Mycobacterium tuberculosis* infection and represent one of the more severe morphological forms of tuberculosis (TB). Several *in vitro* models that mimic human TB granulomas have been reported to decipher complex host-pathogen interactions and discover new prophylactic and therapeutic interventions. They serve as ethical bridge approaches to human studies. However, these models need improvements in generating well-organized granuloma lesions, classic tuberculoma structures, and relevant microenvironments. They are impractical for screening extensive chemical and genetic libraries owing to their low throughput, limited scalability, batch-to-batch variability, and high costs. Here, we describe a ‘mycobacteria-in-spheroid’ co-culture workflow in a standard 96-well plate format that generates a robust 3D cell culture model. This model reproduces key attributes and microenvironments in human tuberculomas and can be scaled up as a high-throughput screening (HTS)-compatible bioplatform. The tuberculoma-like structures generated encompass organized, florid granulomatous lesions and exhibit solid, necrotic, and cavitary morphologies. This model can be developed using freshly isolated human primary cells or a monocytic cell line with virulent mycobacteria. The platform combines the entire workflow from generation to imaging of tuberculoma-like structures *in situ*. It permits the serial quantitation of drug efficacy and monitoring of lesion resolution over several days to weeks following a single treatment. Additionally, we outline a methodology for adopting this workflow for cryo-preservation, enhancing its potential for commercial application. The ease of generation, pliability, cryo-shelf stability, and reproducibility of the bioplatform make it ideal for HTS applications and implementation in the discovery programs of TB and other granulomatous diseases.

## 1. Introduction

Tuberculosis (TB), caused by *Mycobacterium tuberculosis* (*Mtb*), is the world’s leading cause of death from a single infectious agent, with over a million deaths each year (1). A histopathological feature of TB is the formation of dynamic, spatially organized, multicellular clusters called granulomas. These macrophage-rich structures serve as the niche for *Mtb* growth and dissemination while providing the environment where infected macrophages interact with other recruited cells to “wall off” and fight the offending pathogen (2–5). One of the more severe clinical manifestations of TB is the formation of tuberculomas, which conglomerate tuberculous granulomas into well-circumscribed masses, most often in the lungs and brain, resembling cancer tumors in these organs (6–10). Tuberculomas exhibiting a severe morphological form are present in around 5–10% of pulmonary TB patients (11, 12). In pulmonary TB patients, granulomas are highly polymorphic and exhibit a spectrum of structures, including solid, hypoxic, necrotic, and cavitary transformations, and display complex morphologies in the lung sections (13–15). Although spheroid granuloma nodules with a 2–5 mm diameter are common in the lungs, the tuberculoma structure can vary from a few mm to >10 cm in size (11, 14). The cavitary transformation in tuberculomas increases the risk of person-to-person transmission. In addition, it is associated with poor treatment outcomes, relapse, and the likelihood of developing drug resistance in pulmonary TB (16).

Improved and shorter treatment regimens for drug-resistant and drug-susceptible forms of TB are urgently required to meet the goal of TB elimination. Such regimens may arise from a better understanding of the efficacy of new therapeutics within the range of granuloma forms and microenvironments. A growing number of preclinical and clinical studies highlight the importance of designing novel pathogen-targeted and host-directed therapies (HDTs) that readily penetrate and act within the granuloma environment (17–21). Therapeutics that modulate host–*Mtb* interactions in granulomas can be identified using animal models and *in vitro* cell culture systems. Several two-dimensional (2D) and three-dimensional (3D) *in vitro* cell-culture models that mimic nascent TB granulomas have been described in recent years using *Mtb* infection of primary human cells (22–29). Although helpful in screening a limited number of compounds and probing *Mtb*– host interactions (30–32), these models suffer from low throughput, limited tractability, and restricted scalability (33, 34). They form microgranulomas, or small granulomatous aggregates of macrophages, but lack organized lesions and relevant tuberculoma size, structures, and forms.

Consequently, physiological gradients of nutrients, oxygen, pH, and pertinent microenvironments present in heterogeneous TB lesions are either absent in these models or not comprehensively reproduced. Critical features like hypoxia and *Mtb* dormancy observed in some of these 3D models with miniature granulomas primarily result from the encapsulation of macrophages in microspheres or embedding in the extracellular matrix (ECM) rather than the granuloma structures themselves (35–37). The lack of a continuous influx of immune cells in these models makes maintaining dynamic structures and the extended prolongation of experiments challenging. Using animal models for such screening is costly and time-consuming, and it limits the number of compounds that can be screened in a high-containment facility. A high-throughput screening (HTS)-compatible and widely applicable bioplatform that reproduces crucial features and microenvironments in TB lesions, allowing for serial multiparametric readouts of host and pathogen physiology and spatiotemporal existence, is highly desirable.

The protocol described here provides simple workflows for developing an HTS-compatible bioplatform using human monocytes and virulent *Mycobacterium* strains expressing fluorescent proteins, such as bright-red fluorescent protein (tdTomato), for fluorescence intensity and image-based assessment of drug efficacy in 3D cell-culture microplates (**Figure 1**). The ‘mycobacteria-in-spheroid’ co-cultures generated in microwells consistently form 3D tuberculoma-like structures. Since macrophages are cardinal to the core-scaffold formation that shapes host immune responses and therapeutic access in tuberculous granulomas (38), we employed human peripheral blood mononuclear cells (PBMCs) or THP-1 monocytes as a source of macrophages. Methodologies for three different versions of the bioplatform using freshly cultured THP-1 monocytes or purified primary CD14^+^ monocytes from the PBMCs are described (**Figure 2A**–**C**). To our surprise, human immortalized THP-1 monocytes, which possess self-renewal and recruitment capabilities, could develop structurally organized granulomatous lesions in the absence of other myeloid, lymphoid, and non-hematopoietic cells in 3D co-culture.

**Figure 1.**
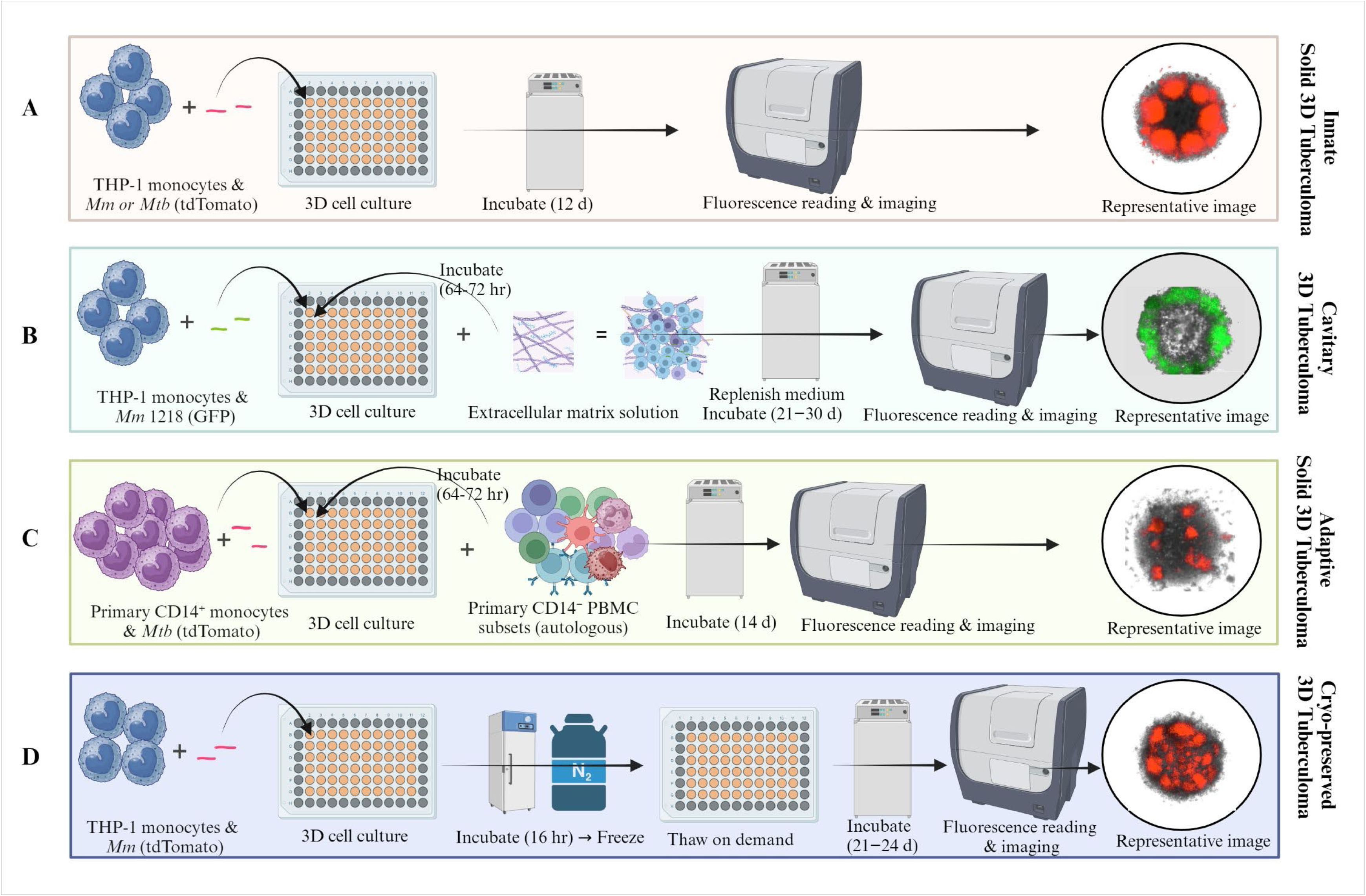
Graphical representation of the ‘mycobacteria-in-spheroid’ co-culture workflows involving human monocytes and fluorescent pathogenic mycobacteria to create a 3D tuberculoma bioplatform. The bio-protocol offers 3D human *in-vitro* granuloma technology with broad applications in tuberculosis. The 3D cell culture workflows generate, (**A**) Innate solid 3D tuberculomas or (**B**) Cavitary 3D tuberculomas using immortalized human monocytes, (**C**) Adaptive solid 3D tuberculomas using human innate and adaptive cell subsets purified from the peripheral blood mononuclear cells (PBMCs), and (**D**) Cryopreserved 3D tuberculomas using immortalized monocytes in a 96-well 3D cell culture microplate. The bioplatform does not require complex materials or specialized equipment to develop and can be coupled with an automated multimode plate reader, live-cell imager, or fluorometer. The protocol primarily spans 2–3 weeks and can be prolonged for several weeks to investigate host-pathogen dynamics or treatment response over an extended period. A versatile, tractable, and high-throughput screening-compatible platform can be used in a BSL-2 or BSL-3 laboratory to screen therapeutic libraries and be preserved for future use. A graphical overview is created in BioRender.com.

**Figure 2.**
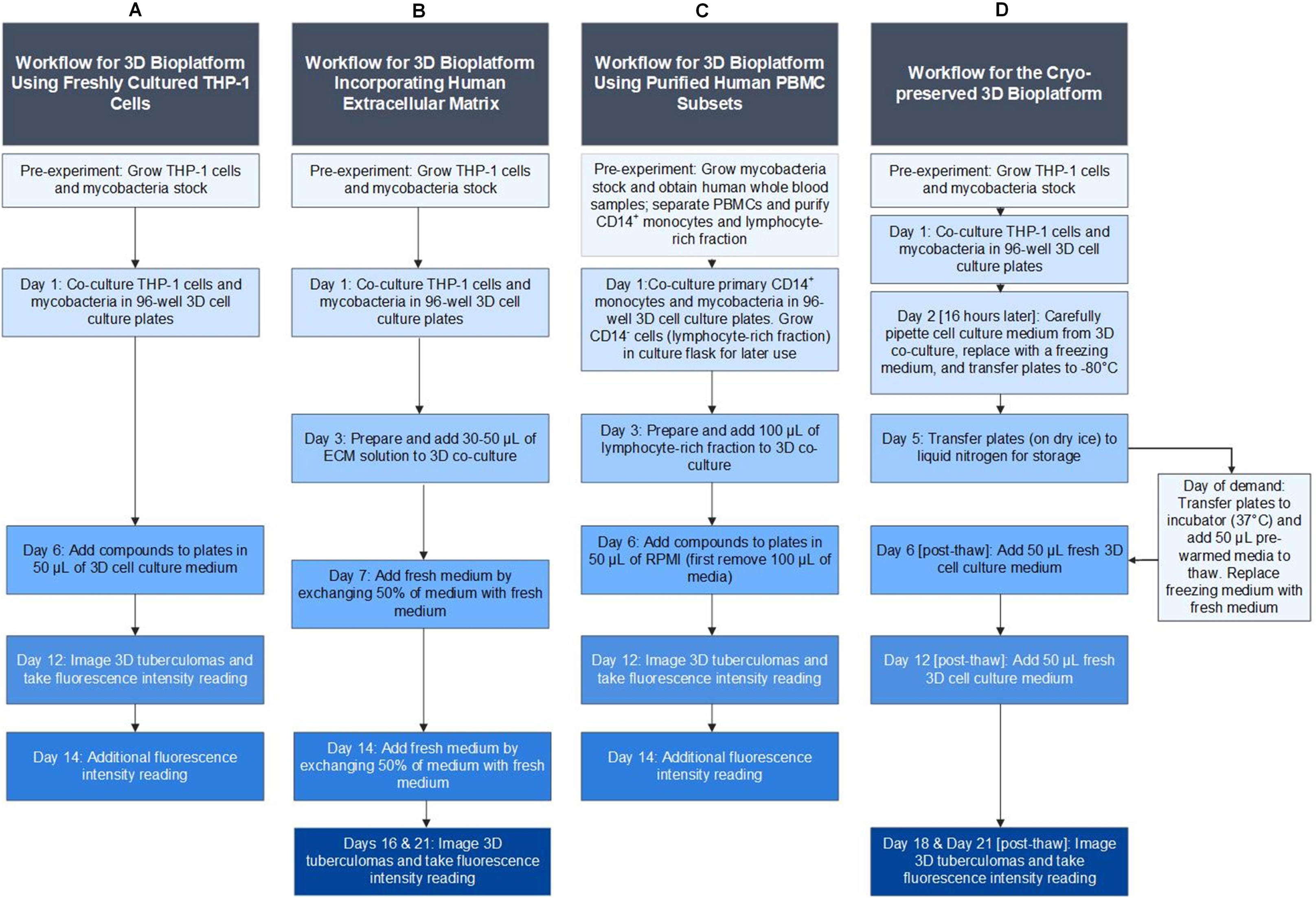
Schematic of the ‘mycobacteria-in-spheroid’ co-culture workflow types in a 96-well 3D cell culture microplate to generate 3D human tuberculoma bioplatform. (**A**) A workflow using freshly cultured immortalized human THP-1 monocytes and pathogenic mycobacteria to generate innate, solid 3D tuberculoma-like structures. (**B**) A modified workflow of THP-1 monocytes and pathogenic mycobacteria incorporating human extracellular matrix to generate 3D tuberculoma-like structures with cavitary features. (**C**) A workflow of purified primary human blood CD14^+^ monocytes and pathogenic mycobacteria that subsequently includes lymphocyte-rich autologous PBMC subsets to generate human donor-specific nascent 3D *in-vitro* tuberculomas with innate and adaptive immune cells. (**D**) A workflow for cryo-stable 3D tuberculoma bioplatform using human THP-1 monocytes and pathogenic mycobacteria that can be frozen for future use and revived on demand to generate solid 3D tuberculomas. 3D, three-dimensional; THP-1 (Tohoku Hospital Pediatrics-1), a monocytic cell line isolated from the peripheral blood of a human leukemia child patient; ECM, extracellular matrix; and PBMC, peripheral blood mononuclear cells.

Furthermore, this 3D spheroid co-culture developed a range of morphological forms that imitate solid, necrotic, and cavitary tuberculomas not described by the available 3D *in-vitro* granuloma models. Other advantages of this platform include ease of development, increased throughput, scalability, pliability, and real-time monitoring of bacterial burden and lesions *in situ*. In addition, we successfully utilized risk group 2 pathogen *M. marinum* (*Mm*) as an alternative to *Mtb* to develop this bioplatform in a BSL-2 laboratory, resulting in a bioplatform comparable to the one created using *Mtb*. *Mm* can cause granulomatous skin infections in humans known as ‘swimming pool or fish tank granulomas’ through contaminated water. In zebrafish, it can develop a human TB-like disease with macrophage-epithelization and tuberculous granulomas (39, 40). The *Mm*– zebrafish model has emerged as a useful *in vivo* platform for studying tuberculous granuloma formation and host-pathogen interactions (38, 41). The workflow with *Mm* can significantly decrease staff hours required during extensive chemical or genetic screening efforts in the BSL-3 laboratory while reducing safety concerns and costs.

Since the successful preservation, maintenance, and long-term storage of the screening bioplatform are highly desirable, we adapted the workflow to develop a cryo-stable version for potential commercialization that can be frozen for future use and revived on demand (**Figure 2D**). The bioplatform and techniques described have the potential to minimize the amount of animal testing in high-containment facilities by improving screening capacity and efficiency, thereby saving time and resources. Even though the system was developed for *Mtb*, we have provided the foundation to allow others to modify this system for other mycobacterial and granulomatous diseases, including TB-associated co-infections and co-morbidities, such as *Mtb* infection with HIV, influenza, or SARS coronavirus-2 and type II diabetes.

## 2. Materials and equipment

### 2.1 Biological materials

1. Human THP-1 cells (American Type Culture Collection (ATCC), catalog number: TIB-202)
2. Human PBMCs from whole blood of healthy, tuberculin skin test-negative donors collected in BD Vacutainer^®^ Cell Preparation Tubes (CPT^TM^) (catalog number: 362761)
3. Fetal bovine serum (FBS), endotoxin-level tested and heat-inactivated (Atlas Biologicals, catalog number: F-0500-A). Store at -10 to -30°C
4. *Mtb* strain H37Rv or Erdman expressing deep-red fluorescent protein tdTomato, *Mm* strain M expressing tdTomato, and *Mm* strain 1218 expressing green fluorescent protein (GFP) ***Critical:*** *The co-culture requires pathogenic mycobacteria (risk group 2 or 3). The attenuated M. bovis BCG strains lack a genomic virulence locus RD1 and are unsuitable*.

### 2.2 Reagents

1. Trypan blue stain (*e.g.*, Invitrogen^TM^, catalog number: T10282). Store at room temperature (RT)
2. Apoptosis/Necrosis kit (Abcam, catalog number: ab176750) (optional). Store at -20°C
3. Invitrogen ImageiT Red Hypoxia Reagent (Thermo Fisher Scientific, catalog number: H10498) (optional). Store at ≤-20°C. Stable for at least 6 months after receipt
4. Acidosis/pHrodo Intracellular pH indicator from Molecular Probes (Thermo Fisher Scientific, catalog number: P35373) (optional). Desiccate and protect from light. Store at 2–8°C
5. Triton X-100 (*e.g.*, Sigma-Aldrich, catalog number: X-100). Store at RT **Caution:** This chemical is harmful if swallowed and causes skin irritation and severe eye damage.
6. Magnetic-assisted cell sorting (MACS) kit LS columns (Miltenyi Biotec, catalog number: 130-042-401). Store at RT
7. *MACS*^®^ CD14 MicroBeads, human CD14^+^ monocyte separation reagent (Miltenyi Biotec, catalog number: 130-050-201). Store at 2–8°C
8. Dimethyl sulfoxide (DMSO), cell-culture grade, and endotoxin level tested (Sigma-Aldrich, catalog numbers: D2650-5×10ML and D5879). Store at RT
9. Polyvinylpyrrolidone (PVP) (MP Biomedicals, catalog number: 102786) (optional). Store at 2–8°C

### 2.3 Media and solutions

1. RPMI 1640 medium with L-glutamine and with or without phenol red (Gibco^TM^, catalog numbers: 11875-093 and 11835-030). Protect from light. Shelf life is 12 months from the date of manufacture. Store at 2–8°C
2. Penicillin-Streptomycin solution, 10,000 units/ml Penicillin and 10,000 µg/ml Streptomycin antibiotics (Gibco^TM^, catalog number: 15140-122). Shelf life is 12 months from the date of manufacture. Store at -5°C to -20°C
3. Sodium pyruvate solution, 100 mM (Gibco^TM^, catalog number: 11360-070). Protect from light. Shelf life is 12 months from the date of manufacture. Store at 2–8°C
4. HEPES buffer, 1M (Gibco^TM^, catalog number: 15630-080). Shelf life is 24 months from the date of manufacture. Store at 2–8°C
5. Mycobacterial liquid growth medium, Middlebrook 7H9 broth with supplements (e.g., prepared in-house, see Recipes. Middlebrook 7H9 broth dehydrated base, BD^TM^, catalog number: 271310; Tween 80, Fisher, catalog number: BP338-500; Middlebrook albumin, dextrose, catalase (ADC) enrichment, BD, catalog number: 212352; and glycerol, Sigma-Aldrich, catalog number: G7893). Store at 2–8°C
6. VitroCol Type 1 human collagen solution (Advanced BioMatrix, catalog number: 5007-20ml). Shelf life is 6 months from the date of receipt. Store at 2–10°C
7. Fibronectin from human plasma, 0.1% solution (Sigma-Aldrich, catalog number: F0895). Store at 2–8°C
8. Sodium hydroxide solution (NaOH) 0.1 M for cell culture (Advanced Biomatrix, catalog number: 5078). Store at RT **Caution:** This chemical causes skin burns and eye damage and is corrosive to metals.
9. Dulbecco’s Phosphate Buffered Saline (PBS), Calcium and Magnesium free, sterile, pH 7.4 (Gibco^TM^, catalog number: 14190-136). Shelf life is 36 months from the date of manufacture. Store at RT
10. Cell culture grade water, sterile (Corning^TM^, catalog number: 25-055-CM). Store at RT
11. Phosphate buffer saline (PBS) 10× without Calcium and Magnesium (Advanced BioMatrix, catalog number: 5076-B). Shelf life is at least 6 months from the date of receipt. Store at RT
12. Red blood cell (RBC) lysis buffer, 1× eBioscience^TM^ (Invitrogen, catalog number: 00-4333-57). Use within 6 months once opened. Store at 2–8°C
13. Ethylenediaminetetraacetic acid (EDTA) solution (0.5M), sterile (Amresco, catalog number: E-177-500ml). Store at RT **Caution:** This chemical can cause irritation to the eyes, skin, and mucous membranes.
14. Leibovitz’s L-15 medium (Gibco^TM^, catalog number: 11415-064) (optional). Protect from light. Shelf life is 36 months from the date of manufacture. Store at 2–8°C
15. Cell growth medium (complete RPMI-1640 medium with antibiotics) (see Recipes)
16. 3D cell culture medium (complete RPMI-1640 medium without antibiotics) (see Recipes)
17. Middlebrook 7H9 broth (see Recipes)
18. Extracellular matrix (ECM) solution (see Recipes)
19. PBMC wash buffer (see Recipes)
20. MACS buffer (see Recipes)
21. 3D cell culture freezing mediums (see Recipes)

### 2.4 Recipes

***Note:*** *The final concentration and amount of each reagent or solution used in the recipes are described below are presented in the **Supplementary Tables 1–9***.

1. Middlebrook 7H9 broth To prepare 1 liter of Middlebrook 7H9 broth, dissolve 4.7 gm of 7H9 broth in 896 ml of distilled water. Add 0.05% (vol/vol) Tween-80 and 0.4% glycerol, mix well, adjust the pH to 6.8–7.0, and filter-sterilize using a 0.2 µm filter assembly. Add 10% ADC aseptically. Store at 2–8°C until use. Determine the sterility of the 7H9 broth by incubating a small volume at 37°C for 48 hr and then at room temperature (15–25°C) for an additional 5 days before use. Stable for at least one month.
2. Cell growth medium To prepare a complete cell growth medium, supplement RPMI 1640 containing L-glutamine (2 mM) with 9.63 % (vol/vol) heat-inactivated FBS, 0.87% (vol/vol) sodium pyruvate solution, 0.87% (vol/vol) HEPES buffer, and 1% (vol/vol) Penicillin-Streptomycin solution and sterile filter using 0.2 µm Nalgene filter assembly. Store at 2–8°C until use. It is stable for at least one month.
3. 3D cell culture medium To prepare a 3D cell culture medium, supplement RPMI 1640 containing L-glutamine (2 mM) with 9.73% (vol/vol) heat-inactivated FBS, 0.88% (vol/vol) sodium pyruvate solution, 0.88% (vol/vol) HEPES buffer, and filter using a 0.2 µm Nalgene filter assembly. Store at 2–8°C until use. It is stable for at least one month.
4. Collagen solution and extracellular matrix (ECM) solution mixture To prepare a collagen solution and an ECM solution mixture, slowly add 1 part of chilled 10× PBS to 8 parts of chilled VitroCol Type 1 human collagen stock solution (3 mg/ml) with gentle swirling. Adjust the mixture’s pH to 7.2–7.6 using sterile 0.1M NaOH. Mix gently by pipetting up and down. Monitor the pH adjustment carefully using pH paper or a pH meter. Adjust the final volume to 10 parts with sterile cell culture-grade water. Maintain the mixture’s temperature at 2–10°C to prevent gelation. Use the solution within 16 hr. Add 40 µl of human fibronectin (0.1%) solution to the 10 ml collagen mixture before use. Mix gently by pipetting up and down.
5. PBMC wash buffer To prepare the PBMC wash buffer required to wash PBMCs after RBC lysis, supplement PBS without calcium and magnesium, pH 7.2–7.4, with 10% (vol/vol) heat-inactivated FBS and 2 mM EDTA. Filter the buffer using a 0.2 µm Nalgene filter assembly. Prepare the buffer on the day of use and keep it cold (2–8°C).
6. MACS magnetic labeling and column elution buffer To prepare the MACS buffer, supplement PBS without calcium and magnesium, pH 7.2–7.4, with 0.5% (vol/vol) heat-inactivated FBS and 2 mM EDTA. Prepare the buffer on the day of use and keep it cold (2–8°C).
7. 3D cell culture freezing media

a. 3D cell culture medium with DMSO To prepare the freezing medium, supplement RPMI 1640-based complete 3D cell culture medium (Recipe 3) with 5% (vol/vol) DMSO. Prepare before use and filter sterilize using a 0.2 µm Nalgene filter assembly.
b. FBS with DMSO (optional) To prepare the freezing medium, supplement a heat-inactivated FBS with 5% (vol/vol) DMSO. Prepare before use and filter-sterilize using a 0.2 µm filter.
c. L15 medium with cryoprotectants (optional) This freezing medium contains equal volumes of serum freezing medium “A” (2×) and DMSO freezing medium “D” (2×). First, add 100 µl of freezing medium “A” to the microwells containing the 3D cell culture, and then add 100 µl of freezing medium “D” before transferring the microplates to a -80°C freezer.
d. Serum freezing medium “A” (2×) To prepare PVP-10× stock, add 10% PVP (wt/vol) to 1× HEPES-buffered saline. Stock 10× HEPES-buffered saline contains 70.7 gm of sodium chloride, 17.0 gm of glucose (dextrose), 2.0 gm of potassium chloride, 19.4 gm of sodium phosphate monobasic 2H_2_O, 47.6 gm of HEPES buffer, and 0.01 gm of phenol red in up to 1000 ml of cell-culture-grade water and 1× HEPES-buffered saline is prepared by adding 900 ml of cell culture grade water in 100 ml of 10× HEPES-buffered saline. To prepare 1000 ml of serum freezing medium “A” (2×), supplement 484 ml of L15 medium with 16 ml of 1M HEPES buffer, 300 ml of heat-inactivated FBS, and 200 ml of PVP-10× stock and filter-sterilize using a 0.2 µm filter.
e. DMSO freezing medium “D” (2×) To prepare 1000 ml of DMSO freezing medium “D” (2×), add 16.02 ml of 1M HEPES buffer and 150.63 ml of DMSO to 833.3 ml of L15 medium, and then filter-sterilize using a 0.2 µm filter. The serum freezing medium “A” and DMSO freezing medium “D”, stored in glass bottles at -20°C, are stable for at least one month.

### 2.5 Laboratory supplies

1. Nalgene® Rapid-Flow™ single-use vacuum filter units, sterile (500 ml, with 0.2 µm membrane) (*e.g*., Thermo Fisher Scientific, catalog number:5660020)
2. Reagent reservoirs, sterile, disposable (*e.g.*, Aquafill, catalog number: S-5501080)
3. Micropipette tips with filters (various volumes), sterile, disposable (*e.g.*, Rainin^TM^, catalog numbers: 30389257, 30389272, 30389276, and 30389274)
4. Serological pipettes (various volumes) (*e.g.*, Pyrex, catalog numbers: 7077-5N and 7077-10N)
5. Cell culture flasks, 75 cm^2^, 150 cm^2^ and 175 cm^2^, sterile (Corning^TM^, catalog numbers: 430720U, 431465, and 431080)
6. Falcon^TM^ conical centrifuge tubes, 15 and 50 ml, sterile (Corning^TM^, Falcon®, catalog numbers: 352097 and 352098)
7. Microcentrifuge tubes (various volumes, sterile) (*e.g.*, GreenTree Scientific, catalog number: T5040G and LabSource, catalog number T56-950)
8. holder or rack (*e.g.*, Fisher Scientific, catalog numbers: 21-200-285, 03-448-17, 21-402-18)
9. Syringe fitted with a needle (25 or 27G), sterile (*e.g.*, Becton Dickinson (BD), catalog number: 309626)
10. 3D cell culture plates (Corning® Spheroid Microplates, catalog number: 4515) ***Note:*** *We tested microplates from several different vendors.* ***Critical:*** *Corning® Ultra-Low Attachment (ULA) Spheroid Microplates were optimal for generating 3D tuberculoma structures described here. This 96-well microplate has an opaque black body that shields each optically clear microwell from well-to-well crosstalk. The round-bottom microwells have a hydrophilic, biologically inert surface. They are coated with a covalently bonded, non-ionic, neutrally charged hydrogel that minimizes activation and cell adhesion to the microwell surface, enabling uniform and reproducible 3D tuberculoma-like structure formation. Another helpful alternative is S-BIO PrimeSurface® 3D Culture Spheroid Plates (S-BIO, catalog number: MS-9096UZ)*
11. Countess™ Cell Counting Chamber Slides (Invitrogen^TM^, catalog number: C-10283) or Neubauer cell counting chamber (*e.g.*, Millipore Sigma, Bright-Line hemocytometer, catalog number: Z359629)
12. Mycobacterial culture bottle (e.g., 490 cm^2^ sterile roller bottle, Corning®, catalog number: 430195)
13. Mycobacterial culture media plates (*e.g.*, Middlebrook 7H10 agar plates without antibiotics and supplemented with oleic acid, albumin, dextrose, and catalase (OADC) enrichment 10% (vol/vol) and glycerol 0.5% (vol/vol) prepared in-house. Middlebrook 7H10 Agar, BD Difco^TM^, catalog number: 262710; Middlebrook OADC, BD, catalog number 212351; and Glycerol, Sigma-Aldrich, catalog number: G7893)
14. Bacterial cell spreaders (*e.g.*, Fisher Scientific, catalog number: 14-665-230)
15. Cell scraper (e.g., Costar^TM^, catalog number: 3010)
16. Petri dish (*e.g.*, Falcon^TM^, catalog: 351029)
17. Assay block 96 well, 2 ml capacity, sterile (Corning^TM^, catalog: 3960)

### 2.6 Equipment

1. Class II Type A2 biological safety cabinet (BSC) (*e.g*., Nuaire, Model: NU-543-600)
2. Micropipettes (various volumes) (*e.g.*, Rainin^TM^, catalog numbers: Pipet-Lite LTS Pipettes L-20XLS, L-200XLS, L-1000XLS, L-5000XLS, and L8-200XLS)
3. Novaspec II spectrophotometer (Amersham Pharmacia Biotech) or equivalent
4. Aerosolve^®^ canisters or equivalent
5. Cell culture incubator, set at 37°C with 5% CO_2_ and >95% humidity (Forma Scientific, Model: 3110)
6. Microcentrifuge for microtubes (Eppendorf, Model: 5430 R)
7. Centrifuge for 15- or 30-ml conical tubes (Eppendorf, Model: 5810 R)
8. Cell counting equipment (Thermo Fisher Scientific, Model: AMQAX1600 Countess II or AMQAX2000 Countess III)
9. pH meter or pH paper (*e.g.*, Fisher brand, catalog number: 13-640-510)
10. Cytation 5 microplate fluorescence reader and cell imager or equivalent with optional CO_2_ source (i.e., CO_2_ controller) (Agilent/BioTek, Model: CYT5MPV with GEN5PRIME and GEN5SPOT) ***Note:*** *Fluorescence plate reader and cell imager other than Cytation 5 can be used*.
11. MidiMACS^TM^ Separator (Miltenyi Biotec, catalog number: 130-042-302) and MACS Multi-Stand (Miltenyi Biotec, catalog number: 130-042-303)
12. Water bath (Precision 180 series)
13. Refrigerator, set at 2 to 8°C
14. Freezer, set at -80°C (range -75 to -85°C)
15. Liquid nitrogen storage for cell culture (temperature range -165 to -196°C)
16. Autoclave

### 2.7 Software

1. Gen5 V3.08 with Spot Counting add-on module (Agilent/BioTek, https://www.biotek.com) ***Note:*** *Gen5 Image+ software controls the operation of the Cytation 5 for photomultiplier tube (PMT)-based microplate reading and automated digital microscopy*.
2. GraphPad Prism V9.3 (GraphPad Prism by Dotmatics, https://www.graphpad.com)

## 3. Methods

### 3.1 ‘Mycobacteria-in-spheroid’ co-cultures and 3D tuberculoma bioplatform

The workflows for generating ‘mycobacteria-in-spheroid’ co-cultures and a freshly engineered or cryopreserved 3D tuberculoma bioplatform are described in detail below in **step-by-step procedures**, including their application for therapeutic screens and the readouts. Essential information regarding mycobacterial cultures and frozen stocks, human primary and immortalized cell cultures, 3D ‘mycobacteria-in-spheroid’ co-cultures, tuberculoma bioplatform, antibiotic or chemical compound treatment, and characterization of microenvironments in the 3D cell cultures using fluorescent probes are described in the workflows. These methods are alternatively summarized in the **Supplementary material** and provided as **Supplementary methods**.

### 3.2 Statistical analysis

The Mann-Whitney test was used to assess the statistical differences between the two groups, while those between three or more groups were measured using the one-way ANOVA and the nonparametric Kruskal–Wallis test with Dunn’s post-hoc test (GraphPad Prism V9.3). A *p* < 0.05 was considered statistically significant, and symbols *, **,***, and **** in the figures indicate *p* < 0.05, < 0.01, < 0.001, and < 0.0001 respectively. The quality of the high-content drug screening assay in the 3D tuberculoma bioplatform using a co-culture of THP-1 monocytes and *Mm* was determined using the Z’ statistic. The Z’-factor describes how well-separated the positive and negative controls are in the HTS assay without the intervention of test compounds. The Z’-factor was calculated by performing the assay in a batch of 8–10 different 3D cell culture microplates as follows:

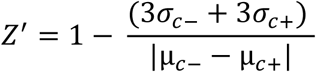

where σ_*c*−_ is the standard deviation of the untreated or DMSO control wells, σσ_*cc*+_ is the standard deviation of the rifampicin or nitazoxanide control wells, µ_*c*−_ is the mean of the untreated or DMSO control wells, and µ_*cc*+_ is the mean of the rifampicin or nitazoxanide control wells. A Z’ value between 0.5 and 1 is considered excellent, a value between 0 and 0.5 is acceptable, and a value less than 0 indicates that the assay is unlikely to be suitable for HTS applications.

### 3.3. Step-by-step procedures

#### A. Workflow for 3D tuberculoma bioplatform using freshly cultured THP-1 monocytes

***Note:*** *This optimized workflow outlines the steps for developing a tuberculoma bioplatform in a 96-well 3D cell culture microplate using a simple co-culture of THP-1 monocytic cells and fluorescent pathogenic mycobacteria for high-content screening (HCS) of potential therapeutics. The typical workflow for screening test compounds, small molecules, or biologics spans 14 days. Depending on the experiment’s goal, the workflow can be extended beyond two weeks if the microwells are replenished with fresh culture medium and THP-1 cells. Using the Mm 1218 strain and carefully exchanging a 50% medium in microwells with a new medium once every seven days, we have successfully cultured the ‘mycobacteria-in-spheroid’ model for up to 50 days*.

***Caution:*** *All work involving the handling of virulent Mtb strains (risk group 3 organisms) should be performed in a BSL-3 laboratory under a Class II BSC, wearing appropriate personal protective equipment (PPE). If the organism of interest is Mm (risk group 2), the experiment is conducted in a BSL-2 laboratory under a Class II BSC, wearing appropriate PPE*.

##### Pre-experiment. Grow bacterial stocks and THP-1 cells

1. Prepare mycobacterial frozen stocks ***Note:*** *We tested Mtb and Mm strains that expressed red fluorescent protein tdTomato for this workflow. In some experiments, Mm strain 1218 GFP was used. The strains used in the study are described in the **Supplementary Methods***.
2. Inoculate 5 ml of 7H9 broth with 100–150 µl of strain (from a strain collection stored at −80°C) in a 25×150 mm glass culture tube (30 ml-capacity) and incubate at 30°C (*Mm*) or 37°C (*Mtb*) for 5–10 days.
3. Measure the inoculum’s optical density at 600 nm (OD_600_). When the OD_600_ reaches 0.6–1.0, add the 5 ml inoculum to 150 ml of 7H9 broth in a 490 cm^2^ sterile culture bottle.
4. Incubate the culture at 30°C (*Mm*) or 37°C (*Mtb*) for 10–12 days, with daily manual shaking, until the OD_600_ reaches 1.0–1.5, as measured by the Novaspec II spectrophotometer.
5. Pellet cultures by centrifugation at 4000× *g* for 30 min at room temperature. Remove the supernatant by decanting or pipetting and then resuspend the pellet in 10 ml of fresh 7H9 broth (Recipe 1).
6. Aliquot the culture at 1.0 ml per cryovial and store at −80°C. Use the 7H9 broth supplemented with appropriate antibiotics to prepare stocks of recombinant fluorescent strains.
7. Thaw an aliquot seven days after freezing and sub-aliquot in the 100 µl working stocks. Determine the viable count on 7H10 agar without antibiotics by incubating at 30°C for 12–15 days (*Mm*) or 37°C for 28–30 days (*Mtb*).
8. Grow THP-1 cells in a cell growth medium (see Recipe 2) ***Note:*** *Follow ATCC’s handling, maintenance, and initial growth procedures*.

a. Seed 175 cm^2^ (T175) tissue culture flask containing pre-warmed (37°C) 50 ml cell growth medium containing antibiotics (Recipe 2) with 4×10^6^ live THP-1 cells from the ongoing cell culture.
b. Place the tissue culture flask in a cell culture incubator set at 37°C with 5% CO2 and >95% humidity to continue the THP-1 cell growth for seven days.

#### Day 1. Development of 3D ‘mycobacteria-in-spheroid’ co-culture

1. Preparing THP-1 cell suspension for 3D cell culture

a. Aliquot 3D cell culture *medium* (see Recipe 3) in 50 ml conical tubes. ***Critical:*** *Verify the sterility of the medium before use*.
b. Prewarm 3D cell culture medium in the cell culture incubator at 37°C for 1 hr.
c. Count THP-1 cells cultured in a growth medium containing antibiotics (Recipe 2) and assess cell viability by the trypan blue dye exclusion method. ***Note:*** *A final concentration of 1×10^6^ live cells/ml is required.* ***Critical:*** *Cell viability must be ≥95% after the last wash. Use fully dissolved trypan blue dye to avoid interference with the automated cell counting caused by the dye precipitates. We used an automatic cell counter, Countess II or III, for cell counting. Cells may be pooled from two or more 150 cm^2^ cell culture flasks cultured in the same batch and subjected to cell count and viability assessment. THP-1 cells cultured in two or more flasks will be required for extensive experiments that require >4 microplates*.
d. Centrifuge cells for 6 min at 200−250× *g* at room temperature (15–20°C) in 50 ml conical tubes and discard the supernatant without disturbing the cell pellet.
e. Resuspend the cell pellet in 15 ml of prewarmed (37°C) 3D cell culture medium (Recipe 3) and gently mix using a pipette. Centrifuge cells again for 6 min at room temperature at 200–250× *g* and discard the supernatant.
f. Repeat this washing step two more times.
g. After the third wash, resuspend the cell pellet in 15 ml of prewarmed (37°C) 3D cell culture medium, mix gently using a pipette, and let the cell suspension rest for 30 min at room temperature. ***Note:*** *This step is incorporated to exchange and remove antibiotics pinocytosed by THP-1 cells during a culture in an antibiotic-containing growth medium in flasks*.
h. Centrifuge cells for 6 min at room temperature at 200–250× *g* and discard the supernatant. Resuspend the cell pellet in 10 ml of prewarmed (37°C) 3D cell culture medium and gently mix using a pipette.
i. Perform the final cell count and assess cell viability using the trypan blue dye exclusion method.
j. Adjust the cell concentration to 1×10^6^ live cells/ml using a 3D cell culture medium. ***Note:*** *Each microplate requires a precise 6 ml of THP-1 cell suspension. Cells are not dispensed in the microwells on the periphery of the plate to avoid an “edge effect.” To mitigate evaporation, the outer wells are filled with sterile 3D cell culture medium or cell culture-grade water*.
2. Preparing *Mtb* H37Rv (tdTomato) or *Mm* M (tdTomato) suspension for 3D cell culture

a. Transport the cryovial containing the frozen stock of *Mtb* H37Rv (tdTomato) or *Mm* M (tdTomato) in an Aerosolve^®^ (transport) canister from a storage freezer to the BSC at the appropriate biosafety level. Place the cryovial in a tube holder in the BSC.
b. Transfer 900 µl of 3D cell culture medium into a microtube containing 100 µl of mycobacterial working stock and mix well.
c. Centrifuge at 5000× *g* for 30 min at 8 to 10°C. When finished, carefully discard the supernatant using a pipette, taking care not to disturb the mycobacterial pellet.
d. Repeat this wash step using a new 1 ml 3D cell culture medium and resuspend the pellet in 1 ml of pre-warmed (37°C) 3D cell culture medium.
e. Pass the mycobacterial suspension through a 25–27G needle fitted to a 1 ml syringe 10 to 20 times to make a single-bacterial-cell suspension. ***Note:*** *Perform this step immediately before infecting THP-1 cells to prevent bacterial aggregation.* ***CAUTION:*** *DO NOT CAP THE NEEDLE ON THE SYRINGE. Place the syringe with a needle in the sharps-disposal container carefully*.
f. Transfer the required amount of mycobacterial single-cell suspension into a sterile reservoir. Dilute mycobacterial suspension using a volume of prewarmed 3D cell culture medium needed to reach the intended multiplicity of infection (MOI) and to make a necessary volume of bacterial inoculum for THP-1 cell infection in microplates. ***Note:*** *Our experiments are routinely performed using five microplates. We typically prepare 35 ml each of THP-1 cell suspension and mycobacterial suspension (infection inoculum) in sterile reservoirs. During the transfer process into microwells, we frequently mixed the inoculum by pipetting up and down to maintain a uniform single-cell suspension and prevent mycobacteria from settling in the reservoir and clumping together. MOI was calculated as the input CFU divided by the number of monocytes added per well. We did not wash monocytes following infection and avoided using aminoglycoside antibiotics to kill extracellular mycobacteria, if any. Pinocytosed aminoglycosides can reach macrophage phagosomes and contribute to cells’ antimicrobial activity* (*42*). ***Critical:*** *The ideal MOI varies with the Mycobacterium species or strains used in the 3D co-culture and is experimentally determined. Refer to the **Results and Data Analysis** sections for the optimal MOI of Mm and Mtb strains used in the 3D co-culture. We used the optimized low MOI of 0.008 (range, 0.006*–*0.012) for co-cultures using Mm M (tdTomato)*.
3. Preparing 3D microplates for 3D cell culture

a. Label Corning^®^ 3D spheroid microplates with plate number, date, experiment, and performer’s name. Mark the peripheral boundary wells. Refer to the **Data Analysis** section for a schematic of the plate used in the compound screening assay.
b. Add 100 µl of THP-1 cell suspension (1×10^5^ live cells) into each microwell except those on the periphery. ***Note****: Use six micropipette tips fitted on a multichannel pipette. Micropipette tips are angled against the wall of the wells during cell suspension dispensing. Therefore, avoid touching the micropipette tips to the bottom of the wells*.
c. Add 100 µl of the mycobacterial suspension to each microwell, except those on the periphery. ***Note****: Control wells with THP-1 cells can be kept without mycobacterial infection, depending on the experiment’s goal. Add 100 µl of 3D cell culture medium to these wells in place of the bacterial suspension*.
d. Mix thoroughly up and down at least three times without touching the microwell bottom to obtain a homogeneous suspension of THP-1 and mycobacteria. Avoid bubble formation during mixing.
e. Fill the peripheral microwells with 250–300 µl of sterile 3D cell culture media or cell culture-grade water using a multichannel micropipette.
f. Place the microplates in a cell culture incubator set at 37°C with 5% CO_2_ and >95% humidity to continue the co-culture experiment and generate 3D ‘mycobacteria-in-spheroid’ structures in ULA round-bottom microwells.
g. Determine the actual CFU count in the 100 µl of mycobacterial suspension by plating dilutions onto Middlebrook 7H10 agar plates. Incubate agar plates at 30°C for *Mm* M (tdTomato) for 12–14 days and 37°C for *Mtb* H37Rv (tdTomato) for 3–4 weeks.

#### Day 6. Addition of test compounds, small molecules, or other therapeutics

1. Preparing and adding drug dilutions for screening in the 3D bioplatform

a. Prepare a dilution of test drugs in a pre-warmed (37 °C) 3D cell culture medium. ***Note:*** *The optimal drug concentration can be experimentally determined. We routinely performed screening experiments using a standard concentration of 20 µM of drug per microwell. In addition, we usually tested six different concentrations, ranging from 20 µM to 0.625 µM, for selected drugs (using 2-fold serial dilution). We performed drug dilutions in 96-well, 2-ml capacity, sterile assay blocks. For screening of large compound libraries, stock compound solutions (10 mM) in DMSO were obtained from the commercial vendors, aliquoted, and stored at -20°C or the recommended temperature. To get a final concentration of 20 µM of test compound per microwell, 10 µl of stock compound (10 mM) was diluted in 990 µl of pre-warmed (37°C) 3D cell culture medium, and 50 µl of this diluted compound solution was added to the microwells containing spheroids in 200 µl of the 3D cell culture medium*.
b. Observe the granulomatous lesion formation in the 3D ‘mycobacteria-in-spheroid’ co-culture using a manual mode in Cytation 5 imager or an inverted microscope (optional). ***Note:*** *Florid granuloma lesions begin to develop between days 5 and 6 in the 3D spheroid infected with the optimized low MOI of Mm M and Mtb H37Rv. The granulomatous lesions grow over time and become structurally organized. We added test drugs on day 6*.
c. Slowly add the diluted drug in a 50 µl volume to the microwells without disturbing the 3D spheroids. Keep a minimum of three technical replicates for individual test drugs and include appropriate positive control (known effective antibiotic or HDT drug) and negative control (medium alone [no-drug added] and drug carrier) wells in each microplate. ***Note:*** *The final volume of the medium in the microwell after drug treatment will be about 250 µl. Negative control wells will receive 50 µl of 3D cell culture medium.* ***Critical:*** *Pipette tips are positioned against the wall of microwells while slowly dispensing the drug solution to minimize disturbance of the formed 3D spheroids and granuloma lesions. Please refer to the **Data analysis** section for more information about positive and negative controls used in our drug screening assay*.

#### Day 12. Fluorescence intensity reading and imaging to assess antitubercular drug efficacy

1. Reading microplates to detect fluorescence intensity as a measure of bacterial burden

a. Place the microplate with lid (without the bottom plate-stand) into the Cytation 5 multi-mode plate reader, previously set to 30°C or 37°C and 5% CO_2._ ***Note:*** *A Cytation 5 combines automated digital widefield microscopy with conventional multi-mode microplate detection to provide phenotypic cellular information and well-based quantitative data. In Gen5 software, ensure that the appropriate plate type (e.g., Corning ULA round well bottom) is selected and the vessel’s bottom elevation is defined. The fluorescence intensity in the co-culture can be measured in the absence of CO_2_ in the Cytation 5 plate reader. We performed in situ fluorescence reading and imaging at 30°C without a CO2 source and controller for Mm co-culture. We tested auto gain and several fixed gain readings in the microplate with 3D spheroids and found the autogain reading suitable for a compound screening assay. For fluorescence intensity readings in the 3D spheroids generated, the ‘top’ reading of the microplate is more optimal than the ‘bottom’ reading. A microplate fluorimeter can serve as a cost-effective alternative to a multimode fluorescence reader. The optimal time to read fluorescence intensity or imaging after drug treatment for the workflow using Mm M (tdTomato) is day 12. The optimal time to read fluorescence intensity or imaging after drug treatment should be experimentally determined for the mycobacterial strain*.
b. Read the tdTomato fluorescence intensity using the fluor-specific fluorescence intensity reading protocol developed, validated, and stored on the computer attached to Cytation 5.
c. To read fluorescence intensity and growth of mycobacteria expressing tdTomato in the Corning 96-well 3D cell culture plate, follow the four steps below.

i. Open the Gen5 software. Next, select the ‘Experiments’ tab on the left panel in the ‘Task Manager’ screen.
ii. On the right-hand side of the screen, choose ‘Create using an existing protocol’ and select the pre-developed and saved tdTomato fluorescence intensity reading protocol. The ‘Task Manager’ screen will be closed. The parameters of the tdTomato fluorescence reading protocol are listed in **Table 1**.
iii. Click the green ‘Read New’ button on the top panel of the Gen5 software. The software will prompt you to save the experiment name and other details in a specific location (directory) on your computer.
iv. After saving the experiment information, the software proceeds to read the fluorescence intensity in the pre-selected microwells of the plate.
d. Export the fluorescence intensity data as an Excel file generated by Gen5 and save it on the computer for data analysis.
2. Automated plate imaging using Cytation 5

a. Place the microplate with lid into Cytation 5 set at 30°C or 37°C and 5% CO_2_.
b. Capture images using the optimized protocol and the parameters listed in **Table 2**. ***Note:*** *The ‘mycobacteria-in-spheroid’ co-culture imaging was carried out to generate quality images of 3D tuberculoma-imitative structures and assess drug efficacy in reducing fluorescent mycobacterial growth and resolution of granuloma lesions. We used a 2.5× objective and a 2×2 image montage to image the entire well. A brightfield imaging channel was used to capture total spheroid images, and a red fluorescence channel (Texas red) was used to capture mycobacteria expressing tdTomato*.
c. To image a 3D cell culture microplate, follow the steps below.

i. Open the Gen5 software. Select the ‘Experiments’ option in the ‘Task Manager’ window.
ii. On the right-hand side of the screen, choose ‘Create using an existing protocol’ and select the pre-developed 3D spheroid imaging protocol. The ‘Task Manager’ screen will be closed. The complete parameters of the imaging and analysis protocol are listed in **Tables 2**–**6**.
iii. Click the ‘Read New’ button on the software’s top panel. The software will prompt you to save the experiment name and details on your computer.
iv. After saving the experiment information, the software will automatically start imaging the preselected microwells in the plate. ***Note:*** *We used montage imaging, image stitching, and Z-stacking in experiment mode in Gen5 to capture 3D tuberculoma images. To capture the entire 3D tuberculoma structure (1800*–*2300 µm in diameter) that spans outside the field of view of the objective, a montage image capture mode was used to capture four image segments (tiles) in the microwell. Individual tiles in the montage are stitched together to make a single, whole image. The ‘auto for stitching’ method was used to capture the montage. The 3D tuberculoma structure exists within a range of Z-planes, and multiple images (slices) must be acquired by moving the objective in the Z-axis (or focal) planes. Therefore, we used a Z-stacking imaging procedure within Gen5 by selecting “Image Z-stack.” Multiple automated image slices were captured below and above the focal plane to ensure that the 3D tuberculomas, cells, and granuloma lesions were imaged at the proper Z-height. After the Z-stacked images are captured, a projection of the Z-stack is created by performing a “Z Projection” step and utilizing the focus stacking algorithm to produce a final composite image. For image stitching and Z-projection, Gen5 Image+ software is required. Each well takes approximately 2 min to complete the imaging process using the described parameters*.
d. Use the parameters listed in **Table 3** to stitch the individual tiles together to generate a single stitched image.
e. For Z-stacking of the individual slices to generate one stacked image, use the parameters in **Table 4**.
3. Cellular analysis of 3D tuberculoma images in Gen5

a. Perform the cellular analysis of projected images using the parameters listed in **Tables 5 and 6**. ***Note:*** *We used cellular analysis to determine the diameter and size of 3D tuberculomas, the number of encompassing granuloma lesions, and the area affected by bacterial growth and lesions in the tuberculoma structures*.

#### Day 13*−*14. Additional fluorescence intensity reading of microplates with 3D culture

***Note:*** *For relatively slow-growing, virulent Mtb strains, such as H37Rv or Erdman (tdTomato), we performed additional fluorescence intensity readings of the co-cultures on day 14. Although fluorescence intensity reading on day 12 provided pertinent information about the efficacy of the test drugs, comparatively better separation of relative fluorescence unit (RFU) readings of test (or positive control) and negative control wells was obtained on day 14 for the workflow using Mtb H37Rv or Erdman. We routinely performed drug screening experiments using five microplates but used up to 12 microplates in some experiments. For experiments employing more than five microplates, imaging can be initiated on day 12, and imaging of the remaining plates can be continued the following day*.

**Table 1.** Fluorescence intensity reading parameters.

| Fluorescence Intensity Reading Parameters |  |
| --- | --- |
| Plate Type | Corning 96-well round-bottom ULA 3D spheroid plate with lid |
| Temperature | 30°C to 37°C |
| Read Method | Fluorescence intensity |
| Read Type | Endpoint |
| Optics Type | Monochromators |
| Fluorophore | tdTomato |
| Excitation | 588 nm |
| Emission | 633 nm |
| Optics Position | Top |
| Gain | Auto |
| Read Speed | Normal |
| Read Height | 7.00 mm |
| Selected Wells | B2 to G11 |

**Table 2.** Automated fluorescence imaging parameters.

| Automated Fluorescence Imaging Parameters |  |  |
| --- | --- | --- |
| Objective | 2.5× FL Zeiss |  |
| Plate Type | Corning 96-well round-bottom ULA 3D spheroid plate with lid |  |
| Temperature | 30°C to 37°C |  |
| Read Method | Image |  |
| Read Type | Endpoint |  |
| Optics Type | Sony Mono CCD Camera |  |
| Brightfield Filter | Total spheroids |  |
| Texas Red Filter | *Mm* M or *Mtb* strain expressing tdTomato |  |
| Fluorescence Excitation | 586 nm |  |
| Fluorescence Emission | 647 nm |  |
| Selected Wells | B2 to G11 |  |
| Channel | Brightfield | Texas Red (586,647) |
| LED | 5 | 10 |
| Integration Time | 137 | 1000 |
| Gain | 1.45 | 15.6 |
| Montage Type | 2 × 2 | 2 × 2 |

**Table 3.** Image stitching parameters.

| 3D Image Stitching Parameters |  |
| --- | --- |
| Registration Channel | Texas Red 586, 647 |
| Montage Size | 2326 × 1718 (7.62 Mb) |
| Fusion Method | Linear Blend |

**Table 4.** Z-stacking parameters for 3D imaging.

| 3D Imaging Z-Stacking Parameters |  |
| --- | --- |
| Focus Method | Fixed focal height (1551 $\mu\text{M}$) |
| Number of Slices | 11<br>(Images taken through the structure) |
| Step Size | 68 $\mu\text{m}$ |
| Sample Thickness | 680 $\mu\text{m}$ |
| Images Below Focus Point | 0 |
| Stacking Method | Focus stacking |
| Channel | Brightfield / Texas Red |
| Size of Max. Filter | 11 pixels |
| Top Slice | 11 |
| Bottom Slice | 1 |

**Table 5.** Cellular analysis parameters (spheroid analysis)

| 3D Spheroid Cellular Analysis Parameters [For Spheroid analysis] |  |
| --- | --- |
| Primary Mask Criteria (On Brightfield channel) |  |
| Threshold | 5000 |
| Background | Light |
| Split Touching Objects | Checked |
| Fill Holes in Masks | Checked |
| Min. Object Size | 1500 $\mu\text{m}$ |
| Max. Object Size | 4000 $\mu\text{m}$ |
| Include Primary Edge Objects | Unchecked |
| Analyze Entire Image | Checked |
| Advanced Detection Options |  |
| Rolling Ball Diameter | Auto |
| Image Smoothing Strength | 12 |
| Evaluate the Background On | 15% of the Lowest Pixels |
| Primary Mask Criteria (On Texas Red channel) |  |
| Background | Dark |
| Measure within a Primary mask | Checked and “Use Primary Mask” |
| Measure within a Secondary mask | Unchecked |
| Count Spots | Checked |
| Min size | 125 $\mu\text{m}$ |
| Max size | 2500 $\mu\text{m}$ |
| Advanced options | Determined by plate fluorescence |
| Calculated Metrics |  |
| Metric of Interest | Cell Count |
|  | Object Size (Spheroid) |
|  | Object Area (Spheroid) |
|  | Object Sum Area (Spheroid) |
|  | Object Spot Count (Lesions) |
|  | Object Sum Spot Count (Lesions) |
|  | Object Spots Area (Lesions) |
|  | Object Spots Sum Area (Lesions) |
|  | Lesion Area Ratio (Percent Area Affected) * |

\*Custom metric of interest: Lesion Area Ratio = Object Spots Sum Area /Object Area

**Table 6.** Cellular analysis parameters (lesion analysis)

| 3D Spheroid Cellular Analysis Parameters [For Lesion analysis] |  |
| --- | --- |
| Primary Mask Criteria (On Texas Red channel) |  |
| Threshold | Auto-45 |
| Background | Dark |
| Split Touching Objects | Checked |
| Fill Holes in Masks | Unchecked |
| Min. Object Size | 150 $\mu\text{m}$ |
| Max. Object Size | 1500 $\mu\text{m}$ |
| Include Primary Edge Objects | Unchecked |
| Analyze Entire Image | Unchecked |
| Advanced Detection Options |  |
| Rolling Ball Diameter | 1000 $\mu\text{m}$ |
| Image Smoothing Strength | 5 |
| Evaluate the Background On | 5% of the Lowest Pixels |
| Calculated Metrics |  |
| Metric of Interest | Cell Count (Lesions) |
|  | Object Size (Lesions) |
|  | Object Area (Lesions) |
|  | Object Sum Area (Lesions) |

## 1. Read the tdTomato fluorescence intensity using the above-described parameters

***Note****: The workflow described above generates a solid tuberculoma-like structure in each microwell with 3D co-culture (See **Results**). The resulting 3D tuberculoma structures are highly uniform in terms of size distribution, bacterial growth, and the granuloma lesions encompassed. The platform, in 96-well format, can be employed as an HCS and imaging assay platform for anti-TB drug discovery (see the **Data processing and analysis** section)*.

### B. Modified workflow for 3D tuberculoma bioplatform incorporating human extracellular matrix

***Note:*** *This workflow details the steps required to develop a bioplatform using a co-culture of THP-1 cells and a pathogenic mycobacterial strain in the presence of physiologically relevant human ECM components. ECM contributes to the architecture of tuberculous granulomas in the human lungs, and its immunopathological destruction leads to cavity formation in TB patients* (*4, 16, 43*)*. In addition, collagen-rich ECM is known to regulate the survival of macrophages in 3D in-vitro granuloma models* (*26*)*. Still, the human 3D in-vitro granuloma model with cavitary transformations has yet to be described. Advances in 3D in-vitro granuloma models that develop cavitary features will enable investigations into the drivers of cavitation and pharmacological interventions to identify HDT drugs that can prevent or treat Mtb-induced tissue destruction and immunopathology. The 3D tuberculoma model described here develops cavity-like features in spheroid structures. The pre-experiment steps for this workflow include growing Mm 1218 GFP stocks and THP-1 cells as described above*.

#### Day 1. Development of 3D ‘mycobacteria-in-spheroid’ co-cultures

1. Prepare the THP-1 cell suspension as described above in Workflow A.
2. Prepare mycobacterial suspension as described in Workflow A. ***Note:*** *We used the Mm 1218 strain expressing GFP to infect freshly grown THP-1. This strain permitted 3D co-culture for a relatively longer duration when 50% of the culture medium in microwells was exchanged with fresh medium once every seven days. We used a low MOI of 0.005 (optimal range, 0.005 to 0.008) to develop tuberculoma-imitative structures with cavity-like features. Optimization of the MOI and the quantity of extracellular matrix added is recommended when using different mycobacterial species or strains. We have yet to investigate the ability of additional Mm or Mtb strains in our collection to develop this feature in 3D cell culture*.
3. Prepare 3D spheroid microplates for 3D co-cultures as described in Workflow A.

#### Day 3. Incorporation of the ECM solution in 3D ‘mycobacteria-in-spheroid’ co-cultures

1. Preparing the ECM solution

a. Add the required quantity of human collagen solution (3mg/ml) into a 15 ml sterile conical tube.
b. Prepare the ECM solution using human collagen and fibronectin solutions as described in Recipe 4. ***Note:*** *We have used a purified human collagen solution, VitroCol*^®^. *VitroCol collagen is naturally secreted from human neonatal fibroblast cells in the in vitro cell culture. The processed and pure form provided by the supplier was used. VitroCol is approximately 97% Type I human collagen, with the remainder comprising Type III collagen. Storage of VitroCol collagen at 2*–*8*°C *is essential.* ***Critical:*** *Do not freeze. While preparing the collagen mixture, gentle but thorough mixing and careful pH monitoring are crucial. Keep the mixture at 4*°C *in the refrigerator or ice to prevent gelation. The mixture can be prepared a day in advance and stored at 2*–*8*°C *overnight in the fridge to allow homogeneous pH adjustment of the solution. The fibronectin solution is added to the mixture at the end*.
2. Adding the ECM solution

a. Add 30*−*50 µl of the ECM solution to each microwell containing the 3D co-culture on day 3. ***Note:*** *Dispense the ECM solution slowly by slanting the micropipette tips against the side wall of the microwells. Control microwells without ECM addition can be kept, depending on the experiment’s objectives. We have investigated the incorporation of different volumes of ECM solution, ranging from 5 to 50 µl, in the 3D co-cultures. The workflow generates a tuberculoma-like structure with cavitary features in each microwell with 3D co-culture and 30−50 µl of ECM incorporation (See **Results**)*.
b. Place the microplates back into the CO_2_ incubator set at 37°C with 5% CO_2_ and >95% humidity and continue the spheroid co-culture by incubating the microplates.

#### Day 7 and day 14. Addition of fresh medium

1. Add fresh 3D cell culture medium (perform as described above)

a. Carefully exchange 50% of the cell culture medium in the microwells with fresh, prewarmed (37°C) medium. ***Note****: Exchange with a new medium is performed for co-cultures grown for more than 2 weeks*.
b. Observe the growth of spheroid co-culture using a Cytation 5 imager or an inverted microscope (optional).
c. Continue the co-culture by incubating the microplates in the CO_2_ incubator.

#### Day 16 and day 21. Imaging and fluorescence intensity reading of microplates with 3D co-culture

1. Take fluorescence intensity reading and image the 3 spheroids

_a._ Place the microplate with lid into Cytation 5 set at 30 or 37°C, with or without 5% CO_2._
b. Read the fluorescence intensity using optimized parameters as described above in **Table 1**, except that the excitation and emission wavelengths for GFP are 488 nm and 510 nm, respectively.
c. Capture images using the optimized protocol and parameters for imaging with a brightfield filter and fluorescence filter as described above in **Table 2**, except for using a GFP filter instead of a Texas red filter.
d. Perform 3D image processing using the 3D montage imaging, stitching, and Z-stacking parameters described above (**Tables 3 and4**). **Note**: *Following the capture of images, a Z-projection of the Z-stack images was performed using the focus stacking algorithm described above in workflow A to create the final composite images*.
e. Perform cellular analysis of 3D projected images using the parameters described above in **Tables 5 and6**.

### C. Workflow for 3D bioplatform using purified human PBMC subsets

***Note:*** *This optimized workflow outlines the sequence of steps required to develop a bioplatform using co-cultures of purified human CD14^+^ blood monocytes and virulent Mtb Erdman (tdTomato) in a 3D cell culture microplate. Profiling of granulomas in Mtb-infected animal models and humans revealed that macrophages constitute a significant portion (>40*– *50%) of granuloma cells. However, the cellular composition can vary with early and late granulomas and granuloma forms* (*44*)*. Primary monocytes that differentiate into macrophages constitute a small portion, approximately 10% of the cells in human PBMC samples. Due to the insufficient numbers of monocytes and macrophages in the PBMC samples, which limit the continuous recruitment and maintenance of 3D in vitro granulomas, whole PBMCs result in the generation of small cellular aggregates rather than organized, dynamic granulomas. Therefore, the purification and enrichment of CD14^+^ monocytes from the PBMCs are critical steps in this workflow to develop the 3D tuberculoma model. In this workflow, 3D ‘mycobacteria-in-spheroid’ co-cultures are developed using MACS column-purified human primary CD14^+^ monocytes that differentiate into macrophages to simulate the formation of the core scaffold in human tuberculomas. The use of virulent Mtb strains is required. A flow-through unbound fraction generated during MACS column CD14^+^ monocyte purification contains CD14 microbead-unlabeled CD14*^−^ *cells (mainly lymphocyte subsets) from the PBMCs. This ‘lymphocyte-rich’ fraction is cultured separately for two days, washed, and added to 3D spheroids generated using purified autologous CD14^+^ monocytes in the 96-well microplates to simulate lymphocytes accumulated on the periphery of human tuberculomas with macrophage-rich centers. Granuloma lesions develop in ‘mycobacteria-in-spheroid’ co-cultures even without adding this ‘lymphocyte-rich’ fraction. This bioplatform, employing innate and adaptive cell subsets purified from the PBMCs, can be used for a confirmatory screening of ‘top hits’ identified from a primary screen in the THP-1 platform. The pre-experiment steps for this workflow include growing Mtb Erdman tdTomato as described above and identifying blood donors*.

#### Day 1. Development of 3D ‘mycobacteria-in-spheroid’ co-culture

1. Separation of the PBMCs from whole human blood ***Note:*** *PBMCs can be isolated from anticoagulated human blood or “buffy coat” by density gradient centrifugation, for example, using Ficoll-Paque™. We used BD Vacutainer® Cell Preparation Tubes (CPT^TM^) for blood collection and PBMC separation. PBMCs were isolated following the manufacturer’s protocol (BD CPT Manual <u>VDP40104-05 pg 1-2 (bdj.co.jp))</u>. We used 200*–*250 ml of blood per human donor to isolate PBMCs and purify CD14^+^ monocytes. The isolated PBMCs and purified CD14^+^ monocytes were enough to develop five microplates.* ***Caution:*** *Human whole blood may contain bloodborne pathogens, including HIV, HBV, and HCV. Blood from the prescreened donors was used, and universal precautions were taken to ensure safety*.
  a. Collect blood into BD Vacutainer® tubes. Gently mix the blood by inverting the tubes 5 to 10 times before centrifugation. ***Note:*** *BD Vacutainer® tubes should be stored at room temperature (18*–*25*°C*) and labeled adequately for human donor identification*.
  b. Centrifuge tubes with a blood sample at room temperature in a horizontal rotor (swing-out head) for 30 min at 1500–1800× *g*.
  c. Carefully pipette out the plasma without disturbing the white blood cell layer below immediately after centrifugation using a pipette. Collect the cell layer containing mononuclear cells into a 50 ml conical tube. The PBMCs from several CPT tubes can be pooled into one conical tube. Centrifuge the tubes at 1200× *g* for 15 min. Remove the supernatant without disturbing the cell pellet.
  d. Lyse the RBCs by resuspending the pellet in 10 ml of 1× RBC lysis buffer for 5 min at room temperature.
  e. Stop the lysis reaction by adding 25 to 30 ml of PBMC wash buffer (see Recipe 5) and gently mixing the cell suspension using a pipette. Centrifuge cells at room temperature at 275–300× *g* for 10 min. Carefully discard the supernatant.
  f. Resuspend the cell pellet in the 20 ml PBMC wash buffer. Repeat the washing step thrice by centrifugation at 200× *g* for 10 min at 15–20 °C. **Note:** These washing steps, performed at low-speed centrifugation, are required to remove platelets.
  g. Resuspend the cell pellet in the PBMC wash buffer (20–25 ml buffer for PBMCs isolated from 200–250 ml of blood). Perform a cell count and assess cell viability using the trypan blue dye exclusion method. ***Critical:*** *Cell viability should be ≥ 95%. Dead cells may bind nonspecifically to MACS MicroBeads during the purification of CD14^+^ monocytes*.
  h. Centrifuge cell suspension at 300× *g* for 10 min. Remove the supernatant. Adjust the final PBMC concentration to 1×10^7^ live cells in 80 µl of MACS buffer (see Recipe 6) to purify CD14^+^ monocytes. ***Critical:*** *A Buffer containing Ca^2+^ or Mg^2+^ is not recommended. Use a cold buffer (2*– *8*°C*) to minimize nonspecific cell labeling*.
2. Purification of CD14^+^ monocytes from PBMCs by positive selection. ***Note:*** *We purified CD14^+^ monocytes from the PBMCs by using the MACS technique and following the manufacturer’s protocol <u>IM0001260.PDF</u> (miltenyibiotec.com)*
  a. Add 20 µl of human CD14 MicroBeads to 10^7^ live PBMCs in 80 µl of MACS buffer and mix well by pipetting to label the CD14^+^ monocytes in PBMCs magnetically. Scale up all the reagents and total volumes accordingly for higher cell numbers. ***Note:*** *MicroBeads are conjugated to monoclonal anti-human CD14 antibodies. Since CD14 lacks a cytoplasmic domain, it is considered that the antibody binding to CD14 does not trigger signal transduction in monocytes. Therefore, it does not affect the phagocytosis of Mtb bacilli*.
  b. Incubate for 15 min at 2–8°C.
  c. Wash cells by adding 1–2 ml of the MACS buffer (Recipe 6) per 10⁷ cells and centrifuge at 300× *g* for 10 min. Remove the supernatant.
  d. Resuspend up to 1×10⁸ cells in 500 µl of the MACS buffer.
  e. Proceed to magnetic separation using LS columns. ***Note:*** *Choose an appropriate MACS Column and MACS Separator according to the number of total cells and CD14^+^ cells. For LS columns, the recommended sample size for leukocytes is 10⁵–10⁸ labeled cells in a total of 1×10⁷ to 2×10⁹ cells*.
  f. Place the LS column in the magnetic field of a MidiMACS separator attached to the MACS MultiStand.
  g. Prepare the column by rinsing it with 3 ml of the MACS buffer (Recipe 6). Label collected effluent as ‘wash.’
  h. Apply the cell suspension onto the column.
  i. Collect unlabeled cells that pass through and wash the column with 3 ml of the MACS buffer and collect total effluent in a 50 ml conical tube. This unlabeled cell fraction contains lymphocytes and mononuclear cells other than CD14^+^ cells. Perform washing steps by adding 3 ml of MACS buffer three times. Label the tube as a ‘lymphocyte-rich’ fraction. ***Note:*** *Add a new buffer during washing steps only when the column reservoir is empty*.
  j. Remove the column from the separator and place it in a new 50 ml conical tube.
  k. Pipette the 5 ml of the MACS buffer onto the column. Immediately flush out the magnetically labeled cells by firmly pushing the plunger into the column. Label the fraction as CD14^+^ monocytes.
  l. Purify CD14^+^ monocytes from all isolated PBMCs from a donor. Centrifuge conical tubes containing purified CD14^+^ monocytes or the ‘lymphocyte-rich’ fraction at 300× *g* for 10 min.
  m. Resuspend the ‘lymphocyte-rich’ fraction in 50 ml of PBMC growth medium (Recipe 2). Transfer cells to a 150 cm^2^ cell culture flask. Culture cells in a cell culture incubator set at 37°C with 5% CO_2_ and >95% humidity for two days.
  n. Resuspend the pellet of purified CD14^+^ monocytes in 10 ml of PBMC growth medium (Recipe 2) and centrifuge at 300× *g* for 10 min. Remove the supernatant.
  o. Wash the purified CD14^+^ monocytes three times by adding 10–15 ml of prewarmed (37°C) 3D cell culture medium (Recipe 3) and centrifuging at 300× *g* for 10 min. Remove the supernatant. Resuspend purified monocytes in 10 ml of 3D cell culture medium.
  p. . Perform cell count and assess cell viability by the trypan blue dye exclusion method.
  q. Adjust the cell concentration of CD14^+^ monocytes to 2×10^6^ live cells/ml using a 3D cell culture medium.
3. Development of 3D ‘mycobacteria-in-spheroid’ co-culture using CD14^+^ monocytes
  a. As described above in Workflow A, develop ‘mycobacteria-in-spheroid’ co-cultures using CD14^+^ monocytes and *Mtb* Erdman (tdTomato) in the 96-well Corning 3D spheroid microplates. ***Note****: Our experiments are routinely performed using 5,000 CFU of Mtb Erdman for the infection of 2×10^5^ live monocytes per microwell (MOI 0.025)*.

#### Day 3. Addition of lymphocytes and other CD14-negative mononuclear cells to the co-culture

1. Preparation and addition of ‘lymphocyte-rich’ cell subsets in the 3D microplate

a. Collect the ‘lymphocyte-rich’ cell fraction from the cell culture flask into a 50 ml conical tube. ***Note****: Use the cell scraper gently to release any adherent cells that have attached to the flask surface during the 2-day culture period*.
b. Centrifuge the cell suspension at room temperature at 300× *g* for 10 min. Remove the supernatant. Resuspend the cell pellet in 15 ml of prewarmed (37°C) 3D cell culture medium (Recipe 3).
c. Wash cells four times by adding 15 ml of prewarmed (37°C) 3D cell culture medium. Centrifuge at 300× *g* for 10 min. Following the third wash, allow the cell suspension to rest for 30 min at room temperature before proceeding to the final centrifugation. After the last wash, resuspend cells in 10 ml of prewarmed (37°C) 3D cell culture medium.
d. Perform a cell count and assess cell viability using the trypan blue dye exclusion method. Then, adjust the cell concentration to 4×10^6^ live cells/ml using a 3D cell culture medium.
e. Add 100 µl of cell suspension (4×10^5^ live cells) into each microwell without disturbing the spheroids formed by CD14^+^ monocytes. Periphery wells in the microplate are not used as described in the above section. ***Note****: Adding the ‘lymphocyte-rich’ cell fraction facilitates the recruitment of additional immune cell types, including those involved in adaptive immunity, in the monocyte- and macrophage-dominant spheroid core*.

#### Day 6. Addition of test compounds or therapeutics

1. Carefully pipette 100 µl of the supernatant culture medium from the microwells without disturbing the 3D spheroids. Next, as described above in Workflow A, add test compounds dissolved in the 50 µl solution.

### Day 12. Fluorescence intensity reading and imaging to assess antitubercular drug efficacy

#### 1. Perform fluorescence intensity reading and automated plate imaging as described in Workflow A for the THP-1 bioplatform

***Note:*** *The workflow generates a complex, solid 3D tuberculoma-like structure incorporating multiple adaptive and immune cell subsets in microwells with co-cultures (See **Results**). The number of granulomatous lesions formed varies from donor to donor. Since primary human blood monocyte-derived macrophages lack the capability for self-renewal, granulomatous lesions may disintegrate relatively quickly after day 12 for some donors*.

### Day 14. Additional fluorescence intensity reading for microplates with *Mtb*-infected 3D culture

## 1. Perform additional fluorescence intensity reading on day 14 as described in Workflow A

### D. Workflow for the cryo-shelf-stable 3D bioplatform

***Note:*** *Cryopreservation is one of the most promising methods of long-term storage of cells and tissues. Therefore, we developed a cryostable bioplatform by freezing 3D co-cultures in situ in microplates at cryogenic temperatures. Three different time points, i.e., 30 min, 16 hr, and 72 hr post-Mm M (tdTomato) infection of THP-1 cells, were tested to determine the optimal time to freeze the microplates with 3D co-cultures. We also investigated three different freezing media (see Recipes 7a, b, and c). The optimized workflow and **Results** below describe the optimal freezing medium and time to freeze, detailing the steps required to develop a shelf-stable bioplatform in the 96-well 3D spheroid microplate. The pre-experiment steps for this workflow include growing Mm M tdTomato stocks and THP-1 cells as described above*.

#### Day 1. Development and cryopreservation of 3D cell cultures

1. Preparing 3D spheroid microplates for co-culture and cryopreservation
2. Prepare THP-1 cell suspension and *Mm* suspension as described above in workflow A.
3. Resuspend the THP-1 cell pellet in the freezing medium (see Recipe 7a) to prepare a 2×10^6^ live cells/ml cell suspension. ***Note:*** *A greater number of THP-1 cells than workflow A is required to compensate for increased cell death during freezing and thawing*.
4. Resuspend the *Mm* M (tdTomato) pellet in the freezing medium (see Recipe 7a) to prepare an 800−1000 CFU/ml suspension.
5. Dispense 100 µl of THP-1 cell suspension (1×10^5^ live cells) into each microwell except those on the periphery.
6. Add 100 µl of the mycobacterial suspension to each microwell except those on the periphery.
7. Mix thoroughly up and down three times without touching the microwell bottom to obtain a homogeneous suspension of THP-1 cells and mycobacteria.
8. Fill the periphery microwells with 250*−*300 µl of 3D cell culture media or sterile cell culture grade water using a multichannel pipette to avoid boundary effect.
9. Keep the microplates at room temperature for up to 30 min if freezing immediately. ***Note:*** *For ‘mycobacteria-in-spheroid’ co-cultures that were intended to be frozen after 16 or 72 hr, we incubated plates at 37°C with 5% CO_2_ and >95% humidity for 16 or 72 hr (see below)*.
10. Place the microplates in cryo-boxes within 30 min post-infection or co-culture and transfer the 3D microplates into a -80°C freezer. **Pause Point.** Keep microplates in a -80°C freezer for 12*−*72 hr.
11. Transfer the microplates to liquid nitrogen at -160°C to -196°C for long-term storage. **Pause Point.** Transport the microplates to the liquid nitrogen storage facility on dry ice. ***Note:*** *For ‘mycobacteria-in-spheroid’ co-cultures intended to be frozen after 16 or 72 hr, we performed the co-culture in 3D cell culture medium in microplates as described in Workflow A. After 16 or 72 hr, the cell culture medium was carefully removed using a micropipette and replaced with a freezing medium. By 72 hr, the mycobacterial infection process in the spheroid is far advanced, with the formation of cell-to-cell junctions and adhesions. We avoided disturbing the early 3D spheroids when adding the freezing medium*.

#### Thawing of 3D co-culture and further culture on demand

1. Thawing of 3D cell co-culture

a. Transport the microplates on dry ice from the liquid nitrogen storage facility to the cell culture laboratory.
b. Thaw frozen co-cultures in a CO_2_ incubator set at 37°C with 5% CO_2_ and >95% humidity for 10 to 15 min. ***Note:*** *We removed microplate lids in a sterile incubator to facilitate quick thawing. Alternatively, microplates with co-culture can be carefully thawed on a platform in a water bath set at 37°C*.
c. Add 50 µl of prewarmed (37°C) 3D cell culture medium per microwell when at least half of the freezing medium ice is melted visibly.
d. Following complete thawing at 37°C after 30 to 35 min, centrifuge the microplates at 430× *g* for 6 min.
e. Carefully pipette out the supernatant freezing medium, being careful not to disturb the cell pellet. Add fresh, 37°C prewarmed medium (200 µl per microwell) using a micropipette. Resuspend the cell pellet by pipetting up and down five times. Centrifuge the microplates at 430–450× *g* for 6 min. Perform this washing step three times to remove the freezing medium. ***Critical:*** *We did not centrifuge the microplates for ‘mycobacteria-in-spheroid’ co-cultures frozen 16 or 72 hr post-infection in microplates, where compact 3D structures (and cell-to-cell junctions) are already formed. Washing steps were performed by carefully adding pre-warmed medium* (37°C) *without disturbing the 3D spheroid structures in microwells*.
f. Resuspend the pellet in 200 µl of prewarmed (37°C) 3D cell culture medium per microwell after the final wash.
g. Place the microplates in a CO_2_ incubator set at 37 °C with 5% CO_2_ and >95% humidity for further incubation and revival of the 3D co-culture and the formation of granuloma lesions.

#### Day 6 and day 12 post-thawing. Addition of fresh 3D cell culture medium

1. Add prewarmed (37°C) 3D cell culture medium (50 µl) without disturbing the 3D spheroids.
2. Return the microplates to the CO_2_ incubator to continue the growth of the 3D cell co-culture.

#### Day 18 and day 21. Fluorescence intensity reading and imaging

1. Perform the fluorescence intensity reading as described in Workflow A.
2. Perform imaging and lesion count as described in Workflow A.

***Note:*** *In a cryopreserved and revived ‘mycobacteria-in-spheroid’ co-culture, lesions develop relatively late compared to the freshly developed co-culture system. In this workflow, the optimal time to treat with investigational compounds is between days 14 and 18 post-revival for assessing drug efficacy*.

## 4. Results

### 4.1 Optimization of infection dose in the 3D co-cultures

We determined the optimal range of bacterial numbers required for infecting monocytes in a 3D co-culture, without over-colonization over 2−3 weeks of incubation in 3D spheroids. The kinetics of mycobacterial proliferation, granulomatous lesion formation, and the resultant pathogenesis in the ‘mycobacteria-in-spheroid’ co-cultures of THP-1 monocytes (1×10^5^) using eight different MOI doses (1 to 6000 CFU) of *Mm* 1218 (GFP) and *Mm* M (tdTomato) were investigated over three weeks in early experiments (**Figure 3A**–**D**). Our findings revealed that a low MOI of 0.006 (600 CFU) resulted in organized granulomatous lesion formation with significantly less spheroid area affected by mycobacterial growth (*p* < 0.05) compared to the standard MOI of 0.12 by days 12 and 16. The optimal MOI for the *Mycobacterium* species and strains used in the 3D co-culture was determined by monitoring the dynamics of bacterial growth and granulomatous lesion formation within the 3D structures, and ascertaining the Z’-factor of the HTS assay performed in the resultant bioplatform (See **Data analysis** and **applications** below). The optimal MOI using *Mm* M (tdTomato) was determined to be 0.008 (i.e., 800 CFU per 10^5^ live THP-1 cells), within a range of 0.006–0.012. For the slow-growing *Mtb* strains H37Rv, Erdman, and Beijing F2 expressing tdTomato, we found that a relatively higher MOI (0.012–0.05) was necessary for achieving an excellent (≥ 0.5) Z’-factor (See original study Sable *et al.* 2025) (45). Specifically, the optimal MOIs were 0.025 for *Mtb* H37Rv (tdTomato) and 0.05 for Erdman (tdTomato). Using a low physiological MOI ensures the complete gathering of bacilli by aggregating THP-1 monocytes during the process of 3D spheroid formation, preventing unwanted colonization outside the spheroid structures over the two weeks, without relying on aminoglycoside antibiotics, which are believed to kill only extracellular bacteria.

**Figure 3.**
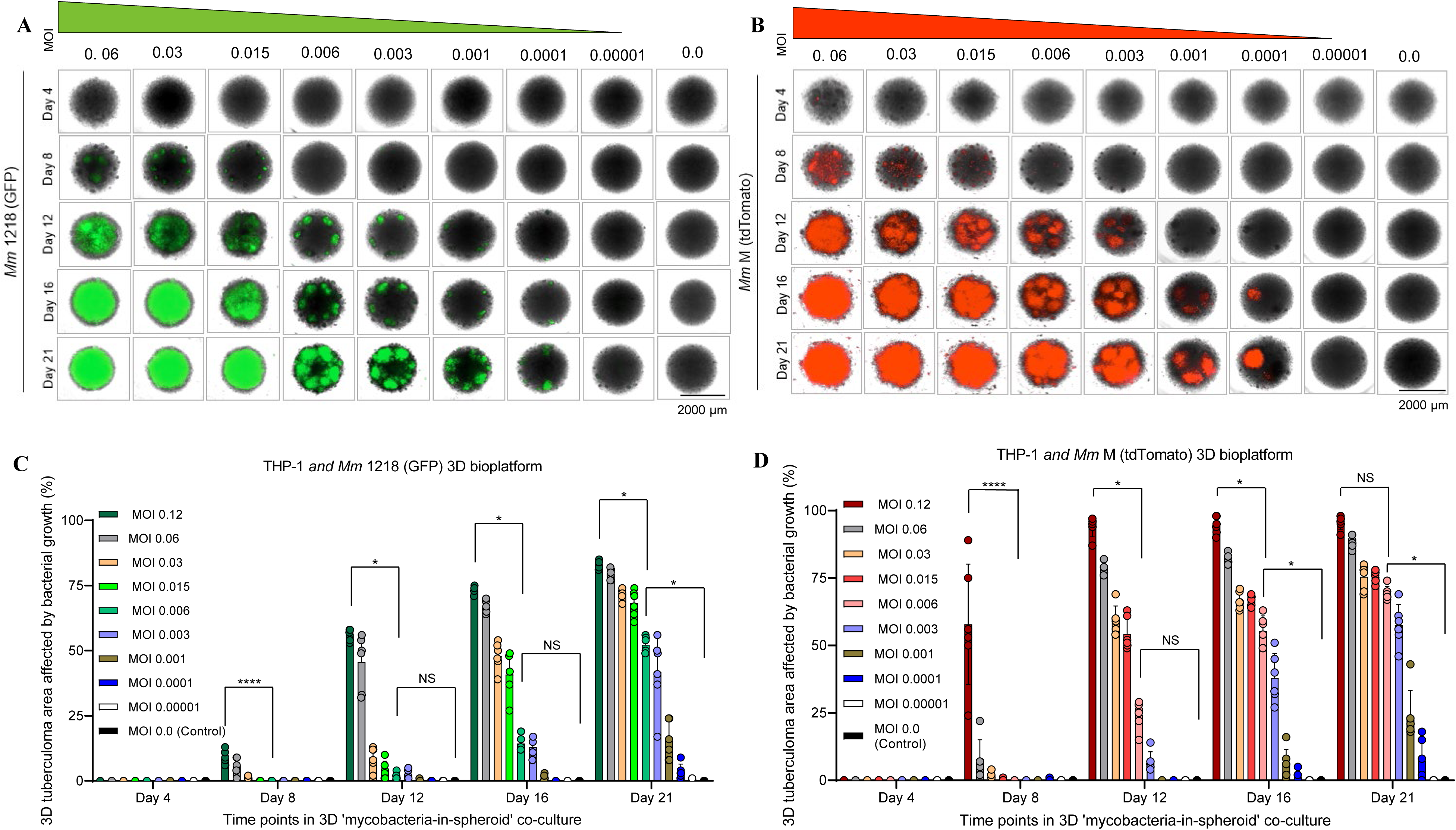
Investigating different multiplicity-of-infection doses of two fluorescent *M. marinum* strains in a ‘mycobacteria-in-spheroid’ co-culture workflow to optimize the 3D tuberculoma bioplatform. (**A**) and (**B**). Kinetics of mycobacteria proliferation, granuloma lesion formation, and the ensuing pathogenesis in ‘mycobacteria-in-spheroid’ co-cultures of THP-1 monocytes (1×10^5^) generated with eight different MOI doses (1 to 6000 CFU) of (**A**) *M. marinum* 1218 (GFP) and (**B**) *M. marinum* M (tdTomato) strain. Representative images are of one of the six 3D spheroids generated using each of the eight MOI doses or no infection controls and captured at five different time points over three weeks of growth from one of the two experiments performed for each strain. Scale 2000 µm. (**C**) and (**D**). Percent area affected by mycobacterial proliferation and granulomatous lesions in the 3D ‘mycobacteria-in-spheroid’ co-cultures generated using THP-1 monocytes (1×10^5^) and nine different increasing MOI doses (1 to 12,000 CFU) of (C) *M. marinum* 1218 (GFP) and (**D**) *M. marinum* M (tdTomato) strain. The circle represents the degree of pathogenesis in one 3D spheroid. The bar represents the average percent area affected by pathogenesis. The data are mean ± SD (n = 6 spheroids) from one of the two experiments. The co-culture using low MOI 0.006 (600 CFU) of *M. marinum* strains results in an organized collection of granuloma lesion formation with <50 and 75% of the 3D tuberculoma area affected by mycobacterial growth by day 12 and 21, respectively. The percentage of affected areas and pathogenesis induced following a low MOI of 0.006 is compared with that caused by the standard MOI of 0.12 and with uninfected controls. * and **** indicate p < 0.05 and < 0.0001, respectively, using the Kruskal–Wallis test followed by Dunn’s post-hoc test. NS, nonsignificant.

### 4.2 Fresh 3D co-cultures of human monocytes and pathogenic mycobacteria

The ‘mycobacteria-in-spheroid’ 3D co-culture workflow, involving freshly cultured human THP-1 monocytes and pathogenic mycobacterial strains *Mm* M, *Mtb* H37Rv, Beijing F2, CDC1551, or Erdman expressing tdTomato (Workflow A), successfully generated solid tuberculoma-like structures in microwells (**Figure 4A**). These 3D tuberculoma-imitative structures encompassed organized granulomatous lesions and developed critical attributes, including hypoxia, necrosis, and acidosis in their cores (**Figure 4B**). In a modified 3D co-culture of THP-1 monocytes and *Mm* 1218 strain expressing GFP and supplemented with human ECM solution (Workflow B), 3D tuberculoma-like structures with cavity-like features were produced (**Figure 4C**). The volume of the ECM solution and the MOI of *Mm* 1218 used influenced the formation of cavitary features. Specifically, 30 µl of ECM solution led to the development of cavity-like features in the 3D co-cultures, but not in the control 3D spheroids. Whereas lower volumes of ECM (<25 µl) or very low MOI (50 CFU) resulted in no cavity formation. In experiments with fluorescent *Mm* strains and human whole PBMCs, which contain fewer than 10% monocytes, small granulomatous aggregates were formed in 3D spheroids (**Figure 4D**) due to the insufficient number of monocyte-derived macrophages to sustain granuloma organization. However, 3D co-cultures of MACS-purified primary CD14^+^ monocytes (2×10^6^/ml) with *Mtb* Erdman (tdTomato) and supplemented with CD14^−^ PBMC subsets (Workflow C) developed larger, organized granuloma lesions, comparable to those formed using immortalized THP-1 monocytes and pathogenic *Mm* or *Mtb* strains. The solid 3D tuberculoma-like structure generated using this workflow thus contains both innate and adaptive immune cell types present in human PBMCs.

**Figure 4.**
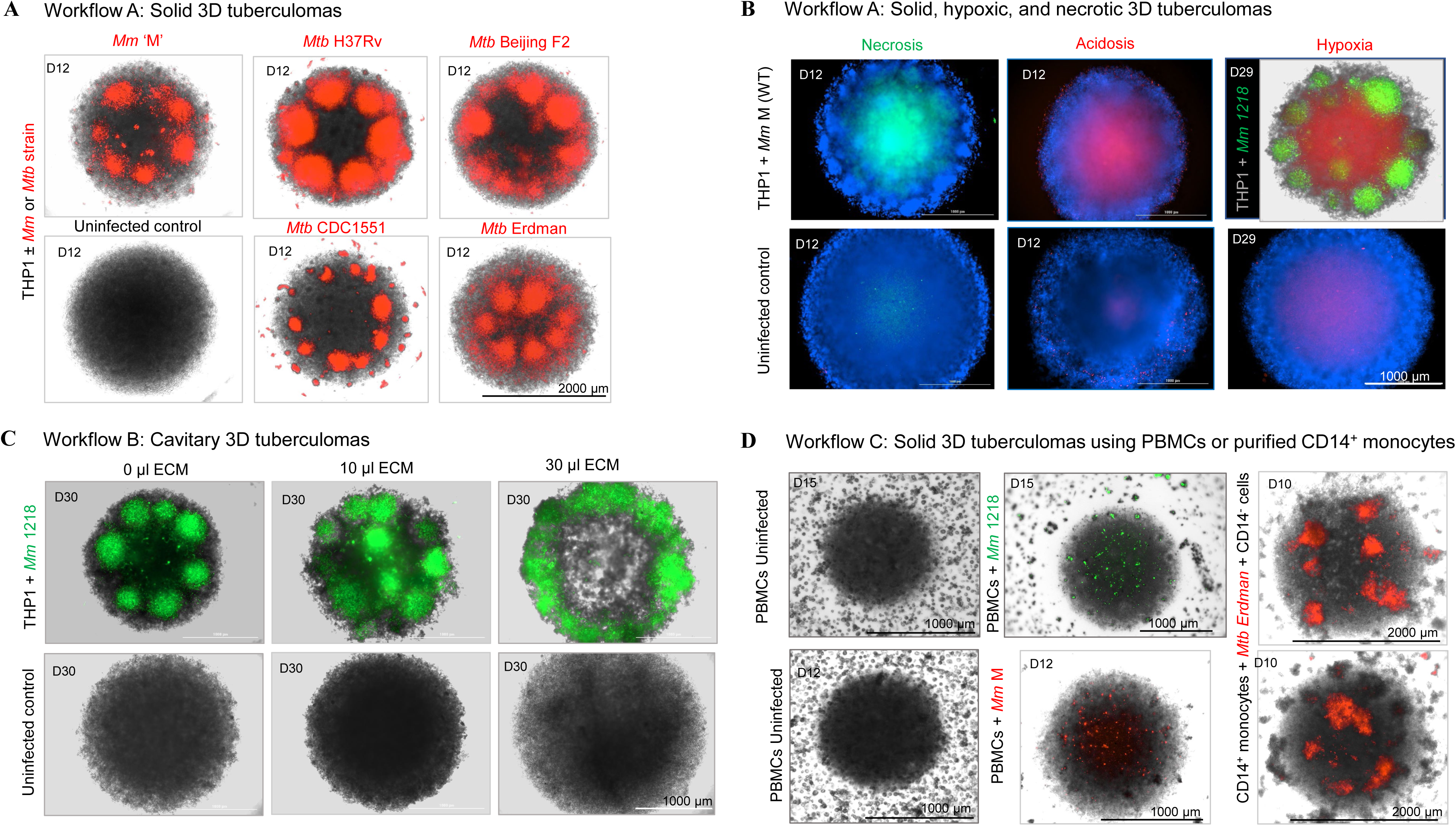
Tuberculoma forms developed using 3D ‘mycobacteria-in-spheroid’ co-culture workflows of freshly cultured human immortalized THP-1 or purified primary blood monocytes and pathogenic mycobacteria. (**A**) Solid tuberculomas generated following 3D co-cultures of freshly cultured THP-1 monocytes and pathogenic, *M. marinum* M, *M. tuberculosis* H37Rv, Beijing F2, CDC1551, or Erdman strain expressing tdTomato fluorescent protein and imaged on day 12. (**B**) Representative images of solid tuberculomas generated following 3D co-cultures of THP-1 monocytes and *M. marinum* wild-type or GFP stained for necrosis (green), acidosis (red), and hypoxia (red) using commercially available fluorogenic probes, as per the manufacturers’ instructions. Representative images of uninfected control spheroids are also shown. Blue counterstain indicates cell nuclei stained using Hoechst 33342 dye. (**C**) 3D tuberculomas generated following a modified 3D co-culture workflow of freshly cultured THP-1 monocytes and pathogenic *M. marinum* 1218 strain expressing GFP and incorporating human ECM comprised of type 1 human collagen and fibronectin (0 µl, 10 µl, or 30 µl). Cavitary features developed in the 3D tuberculomas but not in the control 3D spheroids with 30 µl of ECM. (**D**) Solid tuberculomas generated following 3D co-cultures of total human PBMCs and *M. marinum* 1218 (GFP) or *M. marinum* M (tdTomato) or following 3D co-cultures of purified CD14^+^ monocytes from human blood and *M. tuberculosis* Erdman supplemented with autologous lymphocyte-rich PBMC fraction. While 3D co-cultures of total PBMCs and *M. marinum* developed small granulomatous cellular aggregates, 3D co-cultures of purified CD14^+^ monocytes and *M. tuberculosis* Erdman developed larger, well-defined granuloma lesions. Representative images in (**A**–**D**) are from two or more independent experiments with at least three 3D spheroids.

### 4.3 Cryopreserved 3D co-cultures

We evaluated three freezing media—3D cell-culture medium with 5% DMSO, heat-inactivated FBS with 5% DMSO, and Lebovitz’s L-15 medium with cryoprotective agents—to identify the most effective option for cryopreserving human THP-1 monocyte and *Mm* M (tdTomato) 3D co-cultures (Workflow D). Additionally, we tested three time points (30 min, 16 hr, and 72 hr post-infection) to determine the optimal timing for freezing the microplates with 3D co-cultures. The best results, characterized by organized granulomatous lesion formation after thawing and revival, were achieved when 3D co-cultures were frozen between 30 min and 16 hr post-infection using the 3D cell culture medium with 5% DMSO for cryopreservation (**Figure 5A−F**). In contrast, more extended co-culture periods, exceeding 72 hr post-infection before cryopreservation, resulted in suboptimal outcomes after revival, likely due to the formation of solid 3D structures, cell-to-cell junctions, and nascent lesions.

**Figure 5.**
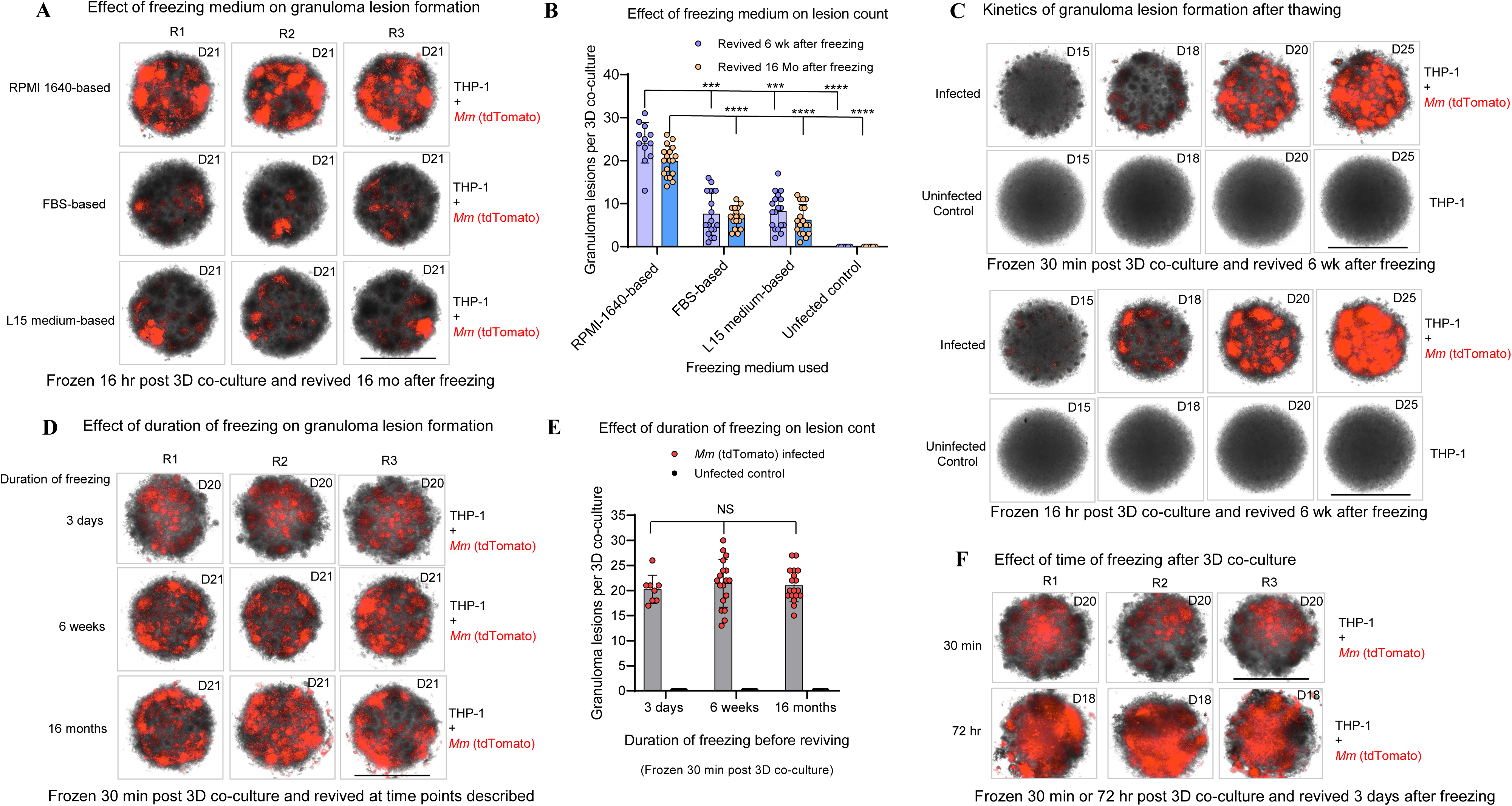
Solid tuberculoma-like structure formation following cryopreservation and revival of 3D co-cultures using THP-1 monocytes and *M. marinum* M. (**A**) and (**B**) Effect of three different freezing media on the granuloma formation in the 3D co-cultures. Significantly more granuloma lesions developed in 3D co-cultures that were frozen 16 hours post-co-culture in the RPMI 1640-based freezing medium compared to the FBS-based or L15 medium-based freezing medium and then revived 6 weeks or 16 months after freezing. (**C**) Kinetics of granuloma lesion formation after thawing frozen 3D co-cultures. A comparable trend in granuloma lesion formation was observed in 3D co-cultures that were frozen 30 minutes or 16 hours post-co-culture in the RPMI 1640-based freezing medium and then revived 6 weeks after freezing. (**D**) and (**E**) Effect of duration of freezing on the granuloma lesion formation. No significant difference in the granuloma count was found in the 3D co-culture frozen in the RPMI 1640-based freezing medium 30 minutes post-co-culture and revived after 3 days, 6 weeks, or 16 months of freezing. (**F**) Effect of time-of-freezing after 3D co-culture on the organized granuloma formation. Numerous well-defined granuloma lesions developed in the 3D co-cultures that were frozen 30 minutes or 16 hours post-co-culture (in (**A**) and (**C**)), compared to 72 hours post-co-culture, in the RPMI 1640-based freezing medium and then revived after 3 days of freezing. Scale 2000 µm. Representative images and data (mean ± SD) are from one to three independent experiments (n = 6–16 spheroids/group). Circles in histograms represent granuloma lesion count per 3D co-culture. ***p < 0.001 and ****p < 0.0001 using the Kruskal–Wallis test with Dunn’s post-hoc test. NS, nonsignificant.

### 4.4 Data processing and analysis

The 3D cell culture workflows using human THP-1 cells or primary CD14^+^ blood monocytes with pathogenic mycobacteria, as described here, were employed in our original study (45). This approach developed 3D structures analogous to solid tuberculomas without the need for ECM embedding, artificial scaffolds, or magnetic levitation. In proof-of-concept experiments, these resulting solid tuberculomas in a 96-well format were used to screen a custom library of known potential HDT chemical compounds. The efficacy of the compounds was evaluated based on their ability to inhibit bacterial burdens and granuloma lesions. Mycobacterial load was determined through fluorescence intensity readings of red fluorescent *Mm* or *Mtb* strains that express tdTomato or by counting CFU on Middlebrook 7H10 agar. This was done after disrupting spheroids and lysing the cells with Triton X-100 (0.1%) for 10 minutes, followed by plating the resulting lysate. Granuloma lesion counts in the 3D structures were obtained through automated imaging and cellular analysis of stitched and Z-stacked images, with manual counts performed by three readers to ensure quality control against Gen5 calculations.

#### 4.4.1 Application in drug screening assay

The bioplatform enables the serial quantitation of drug efficacy *in situ*, in terms of bacterial burden reduction, using a fluorescence plate reader. Additionally, it facilitates the imaging of granulomatous lesion resolution using an automated cell imaging system, as described in the workflows above. The efficacy of test compounds in terms of reduction or increase in bacterial burdens was determined by treating 3D co-cultures with individual test compounds in at least triplicate wells (termed test wells). Wells treated with drug diluent or carrier (i.e., DMSO) and cell culture medium alone without the drug (negative controls) and the antibiotic rifampicin or the experimental HDT drug nitazoxanide (positive controls) were kept in each microplate (see plate map, **Figure 6A**). A pre-defined plate map was used to identify drugs added to microwells. Each microplate contained 3–6 DMSO or carrier-treated wells, 3–6 untreated (no-drug) wells, 3 rifampicin or nitazoxanide-treated wells, and 36 peripheral wells with medium without cells (background) as controls. The normalized reduction of bacterial burden (%) in test compound-treated wells was calculated using the formula below, where Ft is the fluorescence intensity of test compound-treated wells, and Mean Fu is the average fluorescence intensity of untreated (no-drug) or DMSO-treated control wells (n = 3–6 tuberculomas).

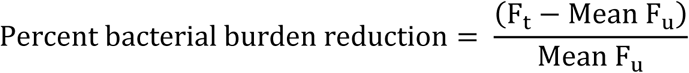

**Figure 6.**
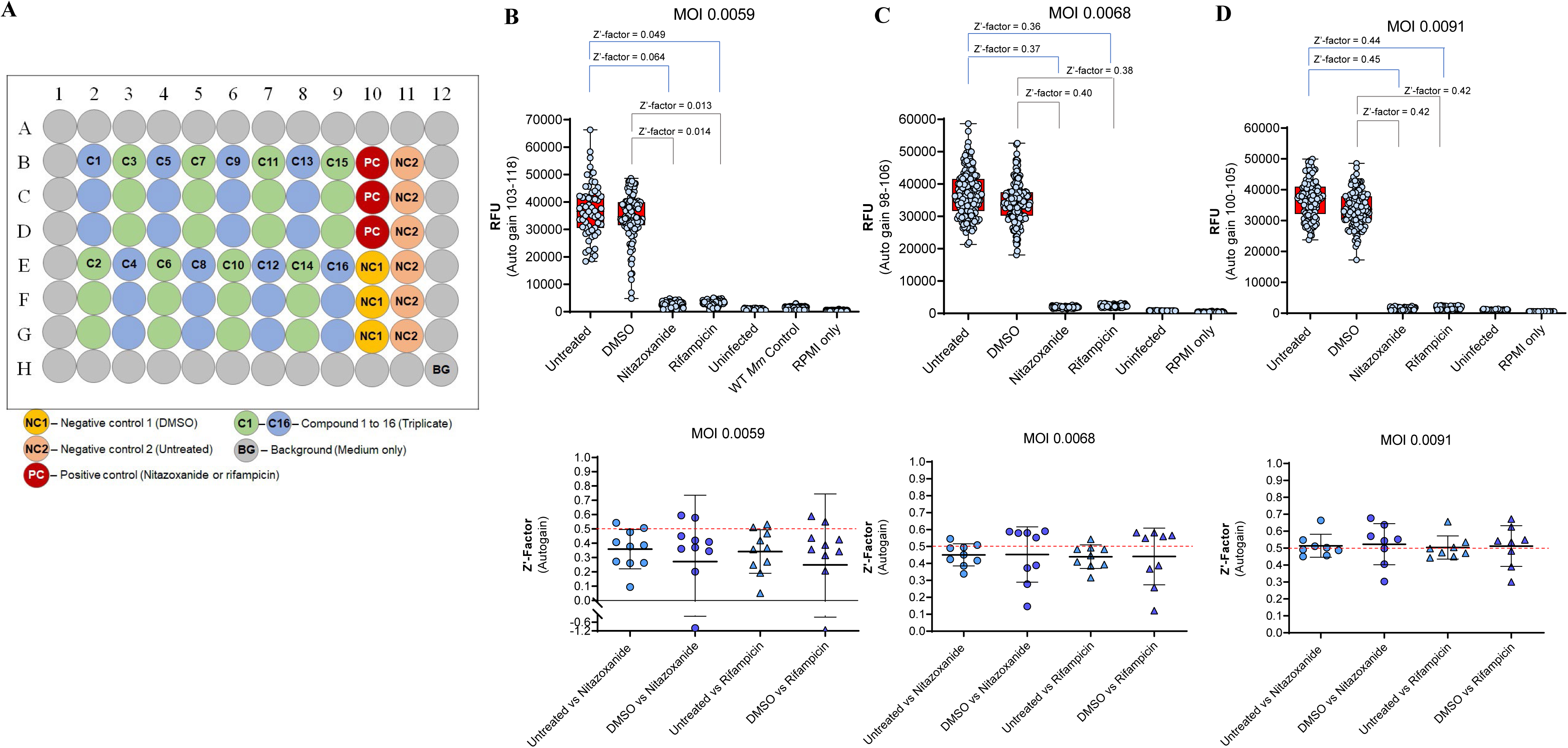
High-throughput screening compatibility and assay quality in 3D tuberculoma bioplatform. (**A**) Plate map showing the typical control and test wells’ location on a 96-well plate. In a primary screen, 16 compounds can be screened in triplicate per plate. (**B**–**D**) Effect of infection dose (MOI) on the Z’-factor of the drug screening assay in the 3D tuberculoma bioplatform developed using THP-1 monocytes and varying infection doses of *M. marinum* M (tdTomato). Three different MOI doses, 0.0059 (**B**), 0.0068 (**C**), and 0.0091 (**D**), were investigated. The upper panels display the Z’ factor in the screening assay, utilizing eight to ten 96-well microplates per MOI experiment, including both negative and positive controls in each plate. Relative fluorescence unit (RFU) data indicating bacterial burdens are presented as box plots with whiskers (minimum to maximum) showing all data points, where a circle symbol represents the RFU level in one well. Fluorescence intensity was measured using auto gain, and the range of auto gain values for microplates used in an experiment is shown. The lower panels show Z’ factors in individual plates of the bioplatform assay, where a symbol represents the Z’ factor in one microplate. The assay using MOI 0.0091 showed an excellent to good assay quality (average Z’ factor ≥ 0.5 shown by the dotted red line).

#### 4.4.2 Quality and suitability of assay for use in HTS

The bioplatform assay reliably detected increased or decreased mycobacterial burdens, granuloma numbers, and lesion size across microwells of several plates per experiment following treatment with test compounds. With rifampicin and nitazoxanide, we obtained a consistent reduction in bacterial burden results (**Figure 6B–D**) with a Z’-factor of >0.4 across microwells and an average Z’-factor of ≥0.5 among several plates (n = 8) when an MOI of 0.009 of *Mm* M (tdTomato) was used in the 3D cell culture microplates (**Figure 6D**), suggesting the acceptable to excellent quality and robustness of the assay using this MOI for HTS applications (refer to statistical analysis section for Z’-statistics). An MOI below 0.006 resulted in a Z’-factor <0 in one out of ten plates, suggesting that this MOI is suboptimal for screening assays. Please also refer to our original study (45) for additional information regarding the assay quality for the *Mtb* H37Rv assay (Z’-factor >0.5) and other data processing and analysis aspects when screening potential therapeutics.

#### 4.4.3 Other applications

Beyond drug screening applications, in the original study (45), we demonstrated the utility of the tuberculoma bioplatform to assess the effects of test compounds simultaneously on host-cell viability, the type of cell death induced, and the potential innate immune mechanisms of action of the test compounds in 3D microenvironments *in situ*. We also demonstrated the utility of this 3D model system in investigating the effects of biologics and biosimilars on the formation and maintenance of granuloma architecture, as well as deciphering early *Mtb*−host interactions in solid or cavitary tuberculoma milieus.

### 4.5 Validation of Protocol

#### 4.5.1 Uniformity and reproducibility of bioplatform

We determined the homogeneity and reproducibility of 3D tuberculoma-like structures formed in the 96-well platform in several microplates and independent experiments. The diameters and areas of 3D tuberculoma-like structures generated in different microplates and batches developed by two performers were measured to determine uniformity and reproducibility. We also analyzed the bacterial burden, the number of granulomatous lesions produced in the 3D spheroids, and the percentage area of the tuberculoma-like structures affected by mycobacterial growth and lesions. The resulting 3D structures exhibited minimal variability (both within and between batches) in size distribution, mycobacterial growth, and granulomas encompassed by individual tuberculoma structures (**Figure 7 A–E**). Their uniformity, reproducibility, and ease of development render them ideal for HTS applications.

**Figure 7.**
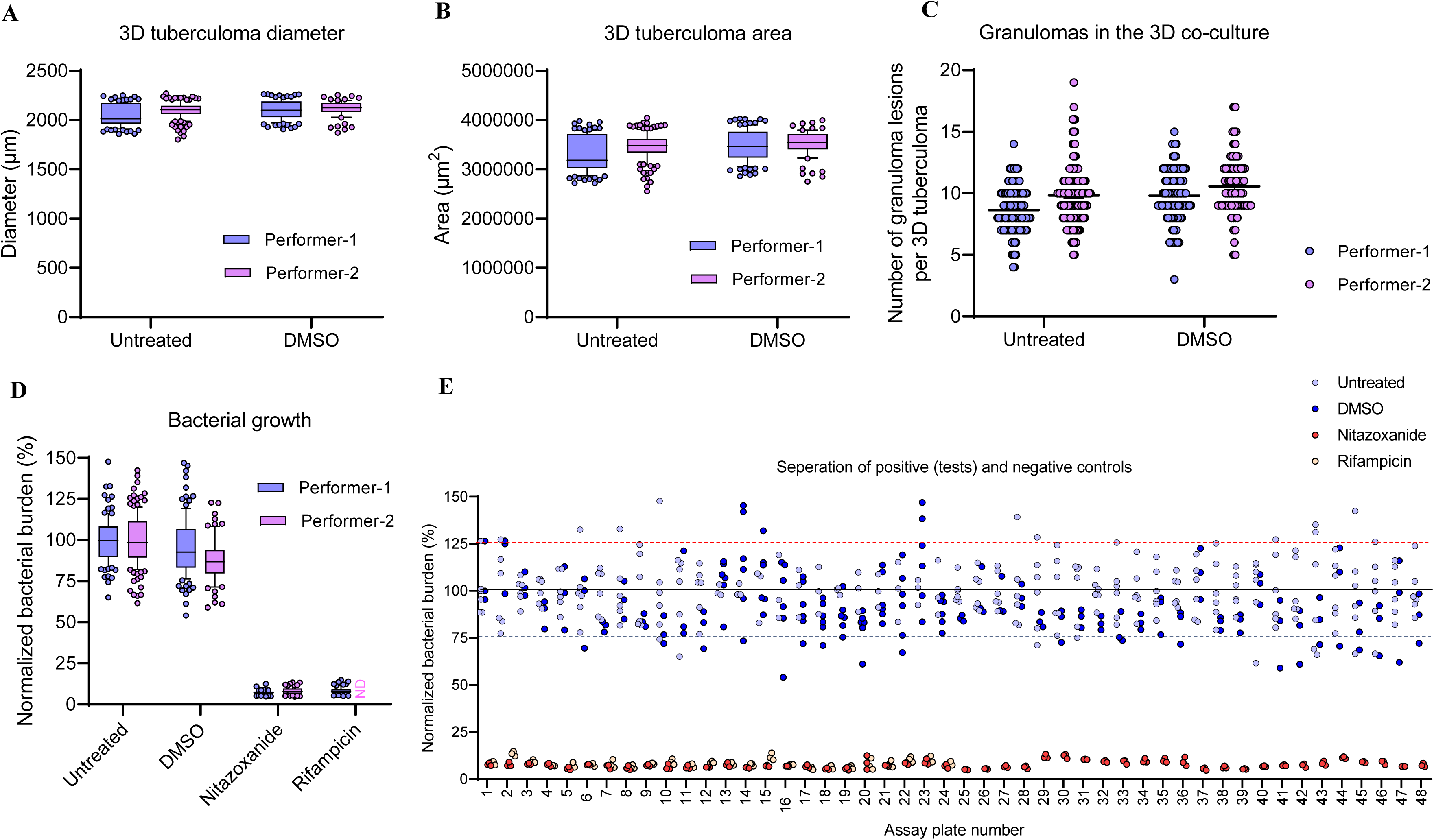
Uniformity and reproducibility of 3D tuberculomas and bioplatform. (**A**−**D**) Consistency in the diameter (**A**), area (**B**), granuloma lesion count (**C**), and bacterial burden (**D**) in the solid 3D tuberculomas developed in 96-well microplates by two separate assay performers using freshly cultured THP-1 monocytes and *M. marinum* M tdTomato. (**E**) Separation of bacterial burden in nitazoxanide or rifampicin-treated (positive controls) and untreated or DMSO-treated (negative controls) 3D tuberculomas. Data in box plots (whiskers: 10–90 percentile) in (**A**–**D**) are from a total of 648 tuberculomas (72–144 tuberculomas per treatment) from 48 different assay plates included in 12 independent experiments (6 experiments per performer) of chemical compound library screenings using the bioplatform assay. Circles represent a response per 3D tuberculoma. The parameters in the 3D tuberculomas developed by the two performers were not significantly different, as determined by the Mann-Whitney test. ND, not done. Bacterial burden data in (**E**) are from 3–6 tuberculomas per treatment from 48 cell culture plates (24 plates per performer) developed for screening potential pathogen-targeted and host-directed compounds using the 3D tuberculoma bioplatform. Rifampicin was not tested in plates 25–48 developed by performer 2.

#### 4.5.2 Other validations

In the preceding sections, we have presented evidence that the developed protocol is highly robust and reproducible. The number of replicates and controls, and the statistical tests used for validation, are described above and in the individual figure legends. For additional details about the characterization of the 3D tuberculoma model developed using primary or immortalized human monocytes, please refer to Figures 2–5 and related Supplementary Figures in the original study (45). For information about how cell culture plate type and culture conditions were selected for 3D *in-vitro* tuberculoma model generation, refer to Figure S2, and for more details about the workflow incorporating human ECM to generate 3D tuberculomas with cavity-like features, see Figure S7 in the original study (45). For additional information about the validation of the utility and the HTS-compatibility of the 3D tuberculoma bioplatform used for screening antimicrobials and HDT compounds within a custom chemical compound library during the proof-of-concept experiments, please refer to Figures 6 and S15–S22 in the original study (45).

## 6. Troubleshooting

Troubleshooting information is provided as **Supplementary Table 10**.

## 7. Discussion

The advanced cell culture systems of pathogenic mycobacteria and human immune cells, resulting in 3D *in-vitro* granuloma models and bioplatforms with HCS capabilities (25–27, 29–31, 33), are highly innovative approaches and have fueled hopes of discovering clinically relevant pathogen-targeting and host-directed therapeutics. However, existing 3D human *in-vitro* granuloma models face challenges regarding ease of generation, throughput, scalability, reproducibility, shelf stability, and screening efficiency. Additional impediments to their utility include technical complexities, difficulties in cell sourcing, limited ability to study granuloma dynamics over time, challenges in retrieving cells for analysis, and limited representation of the microenvironments. A simplistic but high-fidelity bioplatform is highly desirable (33). Yet, the minimum composition and optimal culture conditions for such a facile system for HTS applications, without compromising the TB granuloma formations and tuberculoma attributes, remained unresolved. The observations that the facile 3D co-culture of human primary CD14^+^ monocytes or immortalized THP-1 cells with pathogenic mycobacteria, as minimum essential components, generates crucial features and structures analogous to tuberculomas in the workflows described here are striking. This simplistic ‘mycobacteria-in-spheroid’ 3D co-culture model, in a 96-well format, transforms monocytes into epithelioid macrophages and forms tuberculoma-like structures with a collection of granuloma lesions. This confirms the findings in animal models that the formation of tuberculous granulomas occurs without the contribution of adaptive immunity in the sole setting of innate immunity (2, 41, 46) and provides an HTS-compatible platform with a range of tuberculoma microenvironments for studying host-pathogen interactions and therapeutic screenings.

Despite using spheroid co-cultures to model TB granulomas and generating 3D cellular structures with hypoxia and necrosis, tuberculoma-like features with an organized collection of granulomas or cavitation have not been reported in available models (29, 47, 48). The formation of tuberculoma-like structures and traits in our model required unique spatial features in a culture plate, as well as defined culture conditions and a specific timeframe. 3D spheroid models are ECM-anchorage- or scaffold-independent systems. Macrophages and ancillary cells can contribute to ECM production in 3D cultures. Adding exogenous ECM into the 3D granuloma model prolongs macrophage survival compared to monolayer cultures (26). However, exogenous ECM can reduce granuloma size and limit *Mtb* proliferation (37) and may potentially introduce variability resulting from differences in ECM’s composition and properties. The facile 3D model developed here forms solid tuberculoma-like structures with hypoxic and necrotic cores, without requiring exogenous ECM in the co-cultures, collagen embedding, or polymer encapsulation of macrophages. This approach reduces the system’s complexity while preserving the pliability, scalability, and reproducibility of the bioplatform, ultimately improving throughput.

As described above, the 3D tuberculoma bioplatform has several advantages over existing systems. These include ease of development and use, robustness, self-assembly, cryo-stability, lower costs, increased throughput, and efficiency. The bioplatform also allows real-time monitoring of mycobacterial burdens, granuloma lesions, host cytotoxicity, cavitary transformation, and multiparameter tracking of additional bacterial and host cell physiological attributes *in situ*. The cryopreserved version of the tuberculoma bioplatform can be frozen for future use and revived as needed. The human-relevant model could offer anti-vivisection humane benefits through minimizing animal use in TB research. One of the limitations of the 3D tuberculoma model and workflows described herein is the absence of relevant ancillary cells present in human tuberculous granulomas, such as granulocyte subsets and non-hematopoietic cells, including fibroblasts, epithelial cells, and endothelial cells. Consequently, our model system was unable to investigate the contributions of neutrophils and stromal cells in the formation of necrotizing and caseous tuberculomas and cavitary transformation. To further improve our model, however, we have successfully incorporated these cell types into the THP-1 bioplatform in the form of human cell lines, demonstrating the desired flexibility of our system, while investigating the effects of adding these cell types on granuloma organization in preliminary experiments (data not shown).

Compared to other available 3D human *in-vitro* granuloma models, such as those using polymer encapsulation or matrix embedment, bio-electro-spraying, and magnetic levitation, the methodology described here is straightforward, requiring no specialized instruments or complex materials for development. It provides the tool for investigating heterogeneous granuloma responses to TB vaccines and immunotherapies. It has advantages over the newest 3D systems that facilitate investigations of host-pathogen interactions but lack granuloma formation or 3D tuberculoma features (49, 50). It can be used for siRNA and CRISPRi library screens to identify druggable granuloma host targets. Beyond HTS application for identifying potential HDT compounds and antimicrobials, the bioplatform might allow the development of personalized medicine and therapies using engineered primary human cells, for example, the chimeric antigen receptor (CAR)-macrophages, NK cells, and T cells of patients with difficult-to-treat mycobacterial infections or granulomatous diseases other than TB. The bioplatform can be modified to investigate foreign-body granulomas, non-infectious granulomatous disorders (for example, sarcoidosis, Crohn’s disease, granulomatosis with polyangiitis), and infectious granulomatous diseases, such as those caused by other mycobacteria (e.g., leprosy), bacteria (e.g., brucellosis, cat scratch disease, Whipple’s disease, and *Chromobacterium violaceum* granulomas), fungi (e.g., coccidioidomycosis, cryptococcosis, and histoplasmosis), parasite (e.g., schistosomiasis, leishmaniasis, and toxoplasmosis) and virus (e.g., rubella-induced granulomas).

## Data availability statement

The original contributions presented in the study are included in the article and supplementary material; further inquiries can be directed to the corresponding author.

## Ethics statement

Blood was collected from healthy human donors, and informed consent was obtained from all human subjects. The study followed the protocols and procedures approved by the Institutional Review Board of CDC (IRB protocol number 1652 and project ID number 7294). The mycobacterial species and strains used in the study were obtained from the Laboratory Branch, DTBE, CDC collection. Recombinant strains expressing fluorescent proteins (tdTomato and GFP) were generated using protocols approved by the Institutional Biosafety Committee (IBC protocol numbers 2019.374 and 2020.374). *Mm* M strain, wild-type and recombinant expressing fluorescent tdTomato protein, used in the study, were initially obtained from the collection of Lalita Ramakrishnan Laboratory.

## Author contributions

SBS: Conceptualization, Funding acquisition, Methodology, Investigation, Formal analysis, Data curation, Validation, Project administration, Supervision, Writing – original draft, Writing – review & editing. AK: Methodology, Investigation, Formal analysis, Data curation, Validation, Writing – review & editing. WL: Methodology, Investigation, Formal analysis, Data curation, Validation, Writing – review & editing. JEP: Funding acquisition, Methodology, Project administration, Supervision, Writing – review & editing.

## Acknowledgments

We thank David Tobin, Duke University, Durham, NC, USA, for providing *Mycobacterium marinum* M strains (wild type and recombinant expressing tdTomato protein).

## Generative AI statement

The author(s) declare that no Generative AI was used to create this manuscript.

## Publisher’s note

The findings and conclusions in this publication are those of the authors and do not necessarily represent the official position of the CDC, its affiliated organizations, or those of the publisher, editors, and reviewers. In addition, the trade names used in the study are for identification only and do not constitute an endorsement by the U.S. Department of Health and Human Services, the Public Health Services, or the CDC. Any product that may be evaluated in this article, or claim made by its manufacturer, is not guaranteed or endorsed by the publisher.

This protocol is related to the research article by Sable *et al*. (45), “Identification and characterization of host-directed therapeutics for tuberculosis using a versatile human 3D tuberculoma bioplatform.”

