## Supplemental Methods and Tables for "Methodology for a freshly engineered or cryo-preserved 3D tuberculoma bioplatform for studying tuberculosis biology and high-content screening of therapeutics"

[*]**Correspondence:**

Suraj Sable,

### 1. Supplementary methods

#### 1.1 Mycobacterial cultures and frozen stocks

We used *M. tuberculosis* (*Mtb*) strains, including H37Rv, Erdman, Beijing F2, and CDC 1551, as well as *M. marinum* (*Mm*) strain M expressing the red fluorescent protein tdTomato. The *tdTomato* gene was cloned into the mycobacterial pmsp12 vector using the pTEC27 plasmid. This red fluorescent protein provides optimal brightness with minimal background autofluorescence in 3D spheroid co-cultures. In specific experiments, the *Mm* strain 1218 GFP was used, where the strain *Mm* 1218R expressed GFP utilizing plasmid pFJS8-GFPmut2. Both wild-type and recombinant *Mm* M strains were initially obtained from the laboratory of Dr. Lalita Ramakrishnan. Other mycobacterial strains were obtained from the mycobacterial strain collection of the Laboratory Branch, Division of Tuberculosis Elimination, at Centers for Disease Control and Prevention (CDC), Atlanta. To prepare frozen mycobacterial stocks, fluorescent *Mtb* and *Mm* strains were grown in Middlebrook 7H9 broth (see Recipe 1) with the appropriate selection-antibiotics at either 30°C (*Mm*) or 37°C (*Mtb*). While hygromycin 50 µg/ml was used for strains expressing tdTomato, kanamycin 25 µg/ml was used for a strain expressing GFP. In experiments comparing multiple strains, stocks were prepared simultaneously using the same batch of growth medium and stored at -80°C for one year. After this period, new stocks should be prepared using the original strain collection. A detailed step-by-step procedure for growing mycobacterial stocks is outlined in Workflow A.

#### 1.2 Human primary and immortalized cell cultures

To create 3D cell cultures of immortalized human monocytes, THP-1 cells were grown and maintained according to the guidelines provided by the American Type Culture Collection

(ATCC). The ATCC's handling and growth procedures for cryopreservation, storage in the liquid nitrogen vapor phase, and growing THP-1 monocytes from the frozen stock were followed. The initial growth utilized RPMI-1640-based growth medium containing 10–20% FBS without heat inactivation. THP-1 cells were then grown in a specific cell growth medium (Recipe 2). A detailed step-by-step procedure for cultivating THP-1 monocytes to establish 'mycobacteria-in-spheroid' 3D co-cultures is outlined in Workflow A. For 3D cultures of human primary monocytes, 250 ml of venous blood was collected from healthy human donors in BD Vacutainer Cell Preparation Tubes. Peripheral blood mononuclear cells (PBMCs) were isolated via centrifugation at 1500−1800 *g* for 30 minutes. $CD14^+$ monocytes were purified from PBMCs using CD14 Magnetic-Activated Cell-Sorting (MACS) Microbeads Technology (Miltenyi Biotec). Unlabeled PBMCs were collected as $CD14^-$ cells, which primarily contained lymphocytes and mononuclear cell subsets other than $CD14^+$ monocytes. Purified human mononuclear cells ($CD14^+$ and $CD14^-$) were grown or maintained in the same cell growth medium (Recipe 2) before being used in the 3D 'mycobacteria-in-spheroid' co-cultures. Workflow C outlines the monocyte purification process.

#### 1.3 'Mycobacteria-in-spheroid' co-cultures and 3D tuberculoma bioplatform

Briefly, mycobacterial stocks were washed with 3D cell culture medium (Recipe 3) and centrifuged at 5000 *g*. The resulting mycobacterial suspension was passed through a 25–27G needle fitted to a 1 ml syringe 10–20 times to break bacterial clumps and then diluted in the 3D cell culture medium to achieve the optimal multiplicity of infection (MOI). The optimal MOI of mycobacterial strains expressing fluorescent protein was determined experimentally (See Workflow A and **Results**). The colony-forming unit (CFU) count in the mycobacterial suspension was obtained by plating dilutions onto Middlebrook 7H10 agar plates. For developing 'mycobacteria-in-spheroid' co-cultures and a freshly engineered 3D bioplatform, human immortalized or primary cells were washed with a prewarmed (37 ºC) 3D cell culture medium (Recipe 3) and centrifuged at 250 *g*. A total of $1 \times 10^5$ human THP-1 monocytes or whole PBMCs were seeded in 100 µl of 3D cell culture medium per well in 96-well 3D cell culture plates (Corning® Spheroid Microplates, 4515) and inoculated with 100 µl of the mycobacterial suspension. Cells were mixed by pipetting, and plates were incubated in the cell culture incubator at 37°C with 5% $CO_2$ and 100% humidity for 3D co-culture. Fluorescence readouts and image captures were performed using a Cytation 5 microplate fluorescence reader plus cell imager. In the workflow incorporating ECM, the human ECM mixture was prepared as described in Recipe 4. On day 3, after generating 3D co-cultures of THP-1 monocytes and *Mm* 1218 GFP, 5−50 µl of the ECM solution was carefully added to each microwell. A detailed step-by-step procedure for generating a 3D tuberculoma bioplatform with human ECM is outlined in Workflow B. For 3D co-cultures using purified PBMC subsets, $2 \times 10^5$ purified $CD14^+$ monocytes were infected with *Mtb* Erdman in a 96-well 3D cell culture plate (Corning® Spheroid Microplates, 4515). The cell suspension (200 µl/well) was mixed by pipetting before incubation. On day 3, 100 µl containing $4 \times 10^5$ autologous lymphocyte-rich $CD14^-$ PBMC subsets in the co-culture medium was added to each microwell without disturbing the monocyte-macrophage spheroids. Additionally, to develop 'mycobacteria-in-spheroid' co-cultures and a cryo-shelf-stable 3D bioplatform, we tested three different time points—30 min, 16 hr, and 72 hr post-*Mm* M (tdTomato) infection of THP-1 cells—to determine the optimal time to freeze the microplates with 3D co-cultures. We also investigated three freezing media: 3D cell-culture medium containing 5% DMSO, heat-inactivated FBS containing 5% DMSO, and Lebovitz's L-15 medium containing cryoprotective agents (see Recipes 7a, 7b, and 7c). A detailed, step-by-step procedure for generating a cryo-shelf-stable 3D bioplatform is outlined in Workflow D.

#### 1.4 Antibiotic or chemical compound treatment

The 3D co-cultures of THP-1 monocytes or purified PBMC subsets with *Mm* or *Mtb* strains were established in 96-well 3D cell culture plates, following the procedures outlined in Workflow A or **C**. On day 6, 3D co-cultures developed in 200 µl of medium received treatments in an additional 50 µl of fresh 3D cell culture medium per well. For treatments, antibiotic rifampicin at 1 µg/ml, nitazoxanide, a host-directed and pathogen-targeting compound, at 20 µM, test chemical compounds at 20 µM, DMSO (at equal volume/volume in the medium), or medium alone were used. Mycobacterial growth in co-cultures was monitored in real-time by fluorescence measurements using a Cytation 5 multimode microplate reader and cell imager. The fluorescence readings were taken on days 12 and 14 post-co-culture, and images were captured.

#### 1.5 Characterization of microenvironments using fluorescent probes

The microenvironments in 3D tuberculomas and control spheroids were assessed *in situ* using fluorescent probes and a Cytation 5 cell imager. Necrosis, apoptosis, acidosis, and hypoxia were investigated with commercially available fluorogenic probes, according to the manufacturer's instructions, with some modifications after titrations of the reagents. Hypoxia induction was assessed using the Invitrogen Image-iT$^{TM}$ Red Hypoxia Reagent, a live cell-permeable fluorogenic compound that fluoresces when oxygen levels in the tissue microenvironment reach as low as 5% and reverts when oxygen levels normalize. The reagent has excitation and emission maxima of 490 and 610 nm, respectively. 3D cultures of THP-1 monocytes, with or without the *Mm* strain (M or 1218), were exposed to the reagent at a concentration of 1−10 µM in 3D cell culture medium (Recipe 3). Cultures were incubated in a cell culture incubator at 37°C with 5% CO2, 20% O2, and 100% humidity for 1 hour. The medium was exchanged with fresh 3D cell culture medium, and the cultures were incubated for an additional 3.5 hours before imaging using Cytation 5 with a Texas Red filter.

Apoptosis and necrosis were assessed using the Apoptosis/Necrosis kit (Abcam), which includes Apopxin, a phosphatidylserine (PS) sensor (red, indicating apoptosis), Nuclear Green DCS1, a membrane-impermeable dye that stains the nuclei of damaged or necrotic cells (green, indicating necrosis), and CytoCalcein dye (blue/violet, indicating live cells). After carefully exchanging the culture medium with 100 µl of assay buffer, the spheroids were stained with 200 µl of assay buffer containing 2 µl of Apopxin Deep Red Indicator (100×), 1 µl of Nuclear Green (200×), and 1 µl of CytoCalcein 450 (200×) per well. The spheroids were incubated for 1 hr at room temperature, washed twice with 100 µl of assay buffer, resuspended in fresh assay buffer, and imaged using a Cytation 5 with Texas Red, GFP, and DAPI filters.

For assessing acidosis and intracellular pH, the intracellular pH indicator dye pHrodo$^{TM}$ Red, modified with an acetoxymethyl (AM) ester group, was used. This fluorogenic probe is weakly fluorescent at neutral pH but becomes increasingly fluorescent as pH decreases. The dye traverses the cell membrane and remains within the intracellular space upon cleavage by nonspecific esterases. The excitation and emission spectra of pHrodo are 560 and 585 nm. Briefly, 10 µl of pHrodo$^{TM}$ AM ester, after thawing, was added to 100 µl of Power Load$^{TM}$ Concentrate, a surfactant provided with the kit. This solution was diluted with 10 ml of Live Cell Imaging Solution (LCIS). The cell culture medium of the 3D spheroids was carefully replaced with LCIS and subsequently with pHrodo$^{TM}$ AM ester staining solution. The spheroids were incubated for 30 min at room temperature. The staining solution was replaced with LCIS, and the spheroids were imaged using a Cytation 5.

#### 2. Supplementary Tables

**Supplementary Table 1**: Recipe for Middlebrook 7H9 broth

| Reagent | Final concentration | Amount |
|---|---|---|
| Middlebrook 7H9 broth | 0.47% | 4.7 gm |
| Water (cell culture grade) | 89.1 % | 891 ml |
| Tween 80 | 0.05% | 0.5 ml |
| ADC | 10.0% | 100 ml |
| Glycerol | 0.4% | 4 ml |
| Total | 100% | 1000 ml |

**Supplementary Table 2**: Recipe for cell growth medium

| Reagent | Final concentration | Amount |
|---|---|---|
| RPMI-1640 (1×) | 88.49% | 500 ml |
| FBS (heat-inactivated) | 9.63% | 55 ml |
| HEPES buffer | 0.87% | 5 ml |
| Sodium pyruvate solution | 0.87% | 5 ml |
| Penicillin-Streptomycin solution | 1.00% | 6 ml |
| Total | 100% | 571 ml |

**Supplementary Table 3**: Recipe for 3D cell culture medium

| Reagent | Final concentration | Amount |
|---|---|---|
| RPMI-1640 (1×) | 88.49% | 500 ml |
| FBS (heat-inactivated) | 9.73% | 55 ml |
| HEPES buffer | 0.88% | 5 ml |
| Sodium pyruvate solution | 0.88% | 5 ml |
| Total | 100% | 565 |

**Supplementary Table 4**: Recipe for human collagen extracellular matrix (ECM) solution

| Reagent | Final concentration | Amount |
|---|---|---|
| VitroCol stock solution | 80% | 8 ml |
| PBS (10×) | 10% | 1 ml |
| NaOH solution (0.1M) | N/A | For pH adjustment |
| Water (cell culture grade) | Up to 10% | For adjusting volume (10 parts) |
| Total | 100% | 10 ml |

**Supplementary Table 5**: Recipe for human PBMC wash buffer

| Reagent | Final concentration | Amount |
|---|---|---|
| PBS ($Ca^{2+}$/$Mg^{2+}$ free) (1×) | 89. 6% | 448 ml |
| FBS (heat-inactivated) | 10% | 50 ml |
| EDTA solution (0.5M) | 0.4% | 2 ml |
| Total | 100% | 500 ml |

**Supplementary Table 6**: Recipe for MACS magnetic labeling and column elution buffer

| Reagent | Final concentration | Amount |
|---|---|---|
| PBS ($Ca^{2+}$/$Mg^{2+}$ free) (1×) | 99. 1% | 495.5 ml |
| FBS (heat-inactivated) | 0.5% | 2.5 ml |
| EDTA solution (0.5M) | 0.4% | 2 ml |
| Total | 100% | 500 ml |

**Supplementary Table 7**: Recipe for freezing medium (a), i.e., 3D cell-culture medium with DMSO

| Reagent | Final concentration | Amount |
|---|---|---|
| 3D cell culture medium | 95. 0% | 475.0 ml |
| DMSO | 5.0% | 25 ml |
| Total | 100% | 500 ml |

**Supplementary Table 8**: Recipe for freezing medium (b), i.e., FBS with DMSO

| Reagent | Final concentration | Amount |
|---|---|---|
| FBS | 95. 0% | 47.5 ml |
| DMSO | 5.0% | 2.5 ml |
| Total | 100% | 50 ml |

**Supplementary Table 9**: Recipes for the components of freezing medium (c), i.e., L15 medium with cryoprotectants

1. Serum freezing medium “A” 2×

| Reagent | Final concentration | Amount |
|---|---|---|
| L15 medium (1×) | 48. 4% | 484 ml |
| HEPES buffer (1M) | 1.6% | 16 ml |

| | | |
|---|---|---|
| FBS (heat-inactivated) | 30.0% | 300 ml |
| PVP-10× | 20.0% | 200 ml |
| Total | 100% | 1000 ml |

2. DMSO freezing medium "D" 2×

| Reagent | Final concentration | Amount |
|---|---|---|
| L15 medium | 83. 33% | 833.3 ml |
| HEPES buffer | 1.6% | 16.02 ml |
| DMSO | 15.06% | 150.63 ml |
| Total | 100% | 1000 ml |

**Supplementary Table 10**: Troubleshooting

| Sr. No | Problem | Possible cause(s) | Solution |
|---|---|---|---|
| 1. | 3D tuberculoma structures are not well-organized, and Z'-factor statistics in the tuberculoma bioplatform indicate unsuitability for the HTS assay. | The loss of human monocyte and mycobacterial viability, suboptimal MOI dose, changes in 3D culture plate type, culture conditions, and culture durations can impact the formation of 3D *in vitro* tuberculomas, Z'-factor of the screening assay, and the performance, even though the bioplatform described is very robust. | Optimal bacterial and monocyte viability (>95%), experimentally optimized MOI dose, and standardization of the 3D cell culture parameters are crucial for achieving excellent performance. |
| 2. | Intra-experimental variability in test compounds' host-directed activity and efficacy. | Inaccurate dose or method of compound addition to the plate. | To obtain accurate information about the intracellular and intralesional activity of HDT compounds, we recommend carefully adding drugs to microwells without disrupting the 3D spheroids or rupturing nascent granuloma lesions formed. |
| 3. | 3D tuberculomas of primary monocytes disintegrate quickly before 12 days. | Inter-donor variability or inadequate $CD14^+$ monocyte numbers in 3D co-culture. | For the adaptive immunity model using primary human monocytes and lymphocyte subsets of the PBMCs, 40–60% of purified $CD14^+$ monocytes |

| | | | |
|---|---|---|---|
| | | | (≥2 × $10^5$ monocytes/well) are critical for initiating the 3D co-culture in the absence of differentiated macrophages' self-renewing property, allowing for forming and maintaining organized granulomas over two weeks. |
| 4 | No cavitary feature formation exists in the 3D tuberculoma workflow with ECM. | Culture conditions and parameters are not optimized. | The pH and amount of exogenous human ECM solution added, MOI dose, strain of mycobacteria used for co-culture, and culture conditions and duration can influence the cavitary transformation in 3D tuberculomas. These parameters need careful optimization. |
| 5 | 3D tuberculomas in cryopreserved and revived bioplatform are suboptimal. | 3D cell culture freezing medium and freezing conditions. | The freezing medium and the timing of freezing after 3D co-culture influence the number, structural organization, and kinetics of granuloma lesions formed post-revival in the frozen bioplatform. We recommend carefully standardizing conditions if modifying the parameters described in the cryopreserved bioplatform workflow. |
